# Phase separation of a microtubule plus-end tracking protein into a fluid fractal network

**DOI:** 10.1101/2024.04.19.590270

**Authors:** Mateusz P. Czub, Federico Uliana, Tarik Grubić, Celestino Padeste, Kathryn A. Rosowski, Eric R. Dufresne, Andreas Menzel, Ioannis Vakonakis, Urs Gasser, Michel O. Steinmetz

## Abstract

Microtubule plus-end tracking proteins (+TIPs) are involved in virtually all microtubule-based cellular processes, and it has been recently proposed that they function as liquid condensates. However, the formation process and internal organization of +TIP condensates are poorly understood. Here, we have investigated the phase separation of the CLIP-170 family member Bik1, a key +TIP implicated in budding yeast cell division. We found that Bik1 is a rod-shaped dimer whose conformation is dominated by its central coiled-coil domain. Liquid condensation is accompanied by Bik1 conformational rearrangements, leading to a 2-3-fold rise in interactions between the protein’s folded and disordered domains. In contrast to classical liquids, the supramolecular structure of the Bik1 condensate is heterogeneous, with a fractal structure of protein-rich and protein-free domains. This observation provides structural evidence in support of recent models of biomolecular condensates based on percolation. More broadly, our results provide insights into the structure, dynamic rearrangement, and organization of a complex, multidomain protein in its dilute and condensed phases. Our experimental framework can be extended to other biomolecular condensates, including more intricate +TIP networks.

## Introduction

Microtubule plus-end tracking proteins (+TIPs) are a diverse group of multidomain, often oligomeric, proteins that localize to and track microtubule plus ends, where they form complex and dynamic protein interaction networks.^1^ These +TIP networks are well conserved across species, spanning from yeast to humans. Depending on their capacity to track microtubule ends independently, +TIPs can be categorized into two distinct groups. Autonomous +TIPs, like the members of the end binding (EB) protein family, bind directly to microtubule plus ends and belong to the most conserved and ubiquitous +TIPs.^2^ The second class of TIPs consists of proteins that localize to microtubule ends through direct or indirect interactions with EBs.^3^ Prominent members are the cytoplasmic linker protein 170 (CLIP-170), the adenomatous polyposis tumor suppressor (APC), SLAINs, and the microtubule-actin crosslinking factor (MACF).^4,5^ Notably, +TIPs and their interaction networks play crucial roles in regulating microtubule dynamics and, as such, participate in virtually all microtubule-based cellular processes, including cell division, cell motility, and intracellular signaling.^4^

An example illustrating the involvement of +TIPs in crucial cellular processes is well seen in the budding yeast *Saccharomyces cerevisiae*, where the Kar9-mediated +TIP network drives nuclear positioning during mitosis and mating.^6^ The Kar9 network consists of the core proteins Kar9 (functional homologue of APC, SLAINs, and MACF^7^), Bim1 (EB orthologue), and Bik1 (CLIP-170 orthologue). Specific protein interactions within the Kar9 network have been characterized in the past in large detail,^6–9^ and recent studies have revealed that Kar9, Bim1, and Bik1 exhibit liquid-liquid phase separation (reviewed in ^10–13^) when combined *in vitro*.^14^ In this context, Bik1 has been shown to be the key driver behind the liquid condensation of this protein trio, significantly reducing the critical concentration needed for droplet formation in the mixture.^14^ Importantly, the Kar9-network has been suggested to also form a liquid condensate *in vivo*, dubbed the “+TIP body”, which tracks dynamic microtubule ends and facilitates force transmission between actin cables and microtubule ends during mitosis via the F-actin-directed myosin motor Myo2.^14^ The formation and functioning of the Kar9 +TIP body is predominantly influenced by folded domains, linear motifs present in disordered protein regions, oligomerization domains, and great multivalency and redundancy between interaction partners.^14^ Notably, similar protein elements were reported to mediate the phase separation of +TIPs in other eukaryotic organisms, including EBs and CLIP-170.^15–17^ However, the complexity of +TIP condensates currently hinders our comprehensive understanding of their properties, organization, and modes of regulation.

Bik1 is critically involved in the regulation of microtubule dynamics and participates in nuclear fusion, chromosome disjunction, and nuclear segregation during mitosis in *Saccharomyces cerevisiae.*^18^ Based on the knowledge of CLIP-170’s domain organization and overall shape,^19^ Bik1 is expected to be not a globular but rather an elongated, rod-like protein composed of multiple functional domains that are interspaced by flexible linker regions. Interestingly, Bik1 was recently reported to undergo condensation *in vitro* in reduced ionic strength conditions (change from 500 to 250 mM sodium chloride concentration) without the need of mixing it with other protein partners or crowding agents.^14^ Intriguingly, micrometer-sized Bik1 droplets have an unusually low density and high viscosity compared to other condensates, including FUS.^20^

To deepen our understanding of the formation process and internal organization of +TIP condensates, we here focused on Bik1 as a model system due to its interesting structural and biophysical properties. To this end, we combined computational modeling, microscopy assays, small-angle X-ray scattering (SAXS), and crosslinking mass spectrometry (XL-MS) to study the phase separation of Bik1. Our results provide mechanistic insight into the process of phase separation of a complex dimeric, multidomain protein.

## Results

### Domain organization and phase-separation of Bik1

Until now, only the CAP-Gly domain of Bik1 has been structurally characterized in detail.^8^ To obtain structural information on the 440 amino acid long full-length Bik1, we predicted its structure using AlphaFold.^21^ Based on this computational approach, Bik1 can form an elongated structure (illustrated in **Figure 1A**) with the following domain organization: an N-terminal cytoskeleton-associated protein glycine-rich (CAP-Gly) domain (residues 1-80), a flexible linker region (residues 81-188), a central two-stranded, parallel coiled-coil domain (residues 189-389; its dimeric nature has been assessed previously^22^), and a C-terminal, flexible tail region (residues 390-440) containing a zinc-finger domain (residues 420-431) and the EEY/F-like linear motif QFF (residues 438-440; ETF in CLIP-170).^23,24^ The CAP-Gly domain of Bik1 is important for the proper functioning of the protein and mediates the interaction with the C-terminal EEY/F motif of Bim1^8,25^ and likely with the one of α-tubulin.^26^ Disruption of CAP-Gly’s interaction with EEY/F motifs (Lys46Glu mutation) has been shown to inhibit the accumulation of Bik1 at microtubule plus-ends and, therefore, it obstructs Bik1’s role in regulating microtubule assembly and dynamics.^8,27^

**Figure 1.**
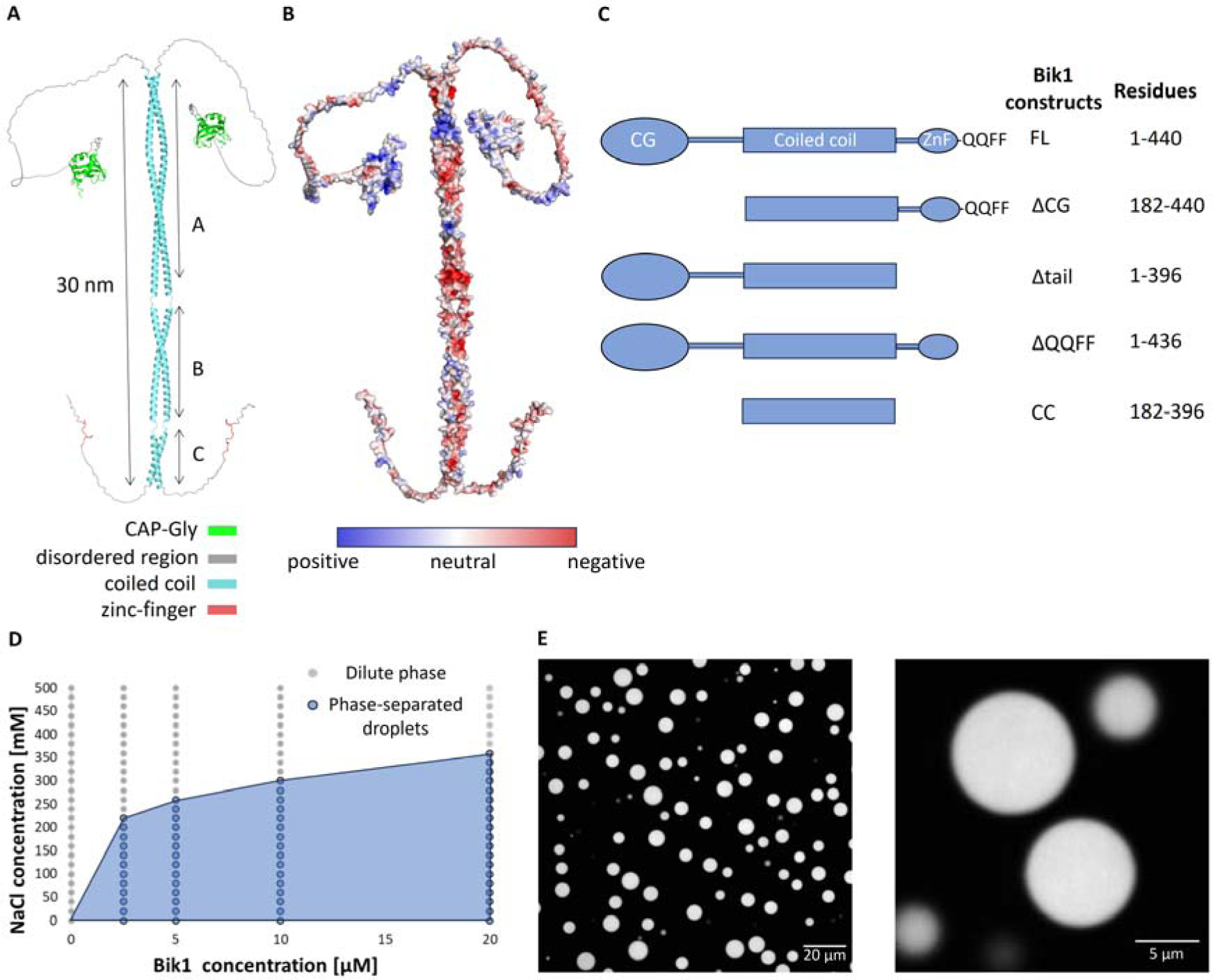
Overview of Bik1 and its variants used in this study. A) One possible structure of the Bik1 dimer as predicted by AlphaFold^21^ (for alternative predicted structures, see **Figure S1**) with color-coded domains. The three predicted segments A, B, and C, of the coiled-coil domain, which are separated by linkers are indicated. B) Charge distribution on Bik1’s surface. Positively charged patches are colored in blue, negatively charged patches are in red, and neutral patches are in light grey. C) Bik1 variants used in this study with their respective designations and residue boundaries. CC, coiled coil; FL, full length, ZnF, zinc finger; QQFF, EEY/F-like motif peptide Gln-Gln-Phe-Phe. D) Phase diagram of N-terminally His-tagged Bik1 FL in 20 mM Tris-HCl, pH 7.4, supplemented with 1 mM DTT and 10% glycerol. E) Confocal fluorescence microscopy images of micrometer-sized droplets observed for 40 µM Bik1 (mixture of 90% Bik1 FL and 10% mNG-Bik1 FL; both proteins are N-terminally His-tagged) in a buffer consisting of 20 mM Tris-HCl, pH 7.4, supplemented with 250 mM NaCl and 2% glycerol. View from the top (left) and a mid-plane slice (right).

While alternative predicted, dimeric structures of Bik1 (**Figure S1**) broadly align with the domain boundaries, they significantly diverge in terms of overall molecular shapes. Notably, the coiled-coil domain is predicted by AlphaFold^21^ and DeepCoil^28^ to consist of three segments (segment A: residues 189-297; segment B: 302-357; segment C: 363-389) that are interrupted by discontinuities in the heptad-repeat sequence of the coiled coil (residues 298-301 between segments A and B (A-B linker), and residues 358-362 between segments B and C (B-C linker); **Figures 1A** and **S1B**). These coiled-coil segment linkers act as hinges that allow the protein to adopt either an elongated or “V-shaped”, folded-back conformations (**Figure S1A**). The CAP-Gly domain is connected to the coiled coil via a flexible linker and as shown in the various AlphaFold predicted structures, can be located at different distances from the coiled-coil domain. The CAP-Gly domain surface is predominantly positively charged, while other domains and regions of Bik1 contain several positively and negatively charged patches along the elongated structure of the protein (**Figure 1B**).

To assess the phase separation of full-length, N-terminally His-tagged Bik1, we recombinantly expressed and purified the protein from bacteria (Bik1 FL; **Figures 1C** and **S2C**). The circular dichroism (CD) spectrum and thermal unfolding profile recorded at 222 nm for Bik1 FL were characteristic of well-folded, α-helical coiled-coil structures (**Figure S2**).^29^ As shown in **Figure 1C**, the phase separation of Bik1 FL depends on both protein and salt concentrations (**Figure 1** Error! Reference source not found.**D**) and gives rise to the formation of micrometer-sized droplets that have been previously observed to merge over time (liquid droplet-like behavior; **Figure S3**).^14^ Notably, the presence of the N-terminal His-tag did not affect the protein’s behavior or its ability to undergo phase separation under the conditions tested (**Figure S3**). To investigate the protein distribution within Bik1 FL droplets, we performed confocal fluorescence microscopy experiments with an N-terminal His-mNeonGreen fusion version of Bik1 FL (mNG-Bik1 FL). These fluorescence microscopy experiments suggest that the distribution of Bik1 dimers is uniform throughout the droplet; the fluorescence of Bik1 is over 60 times brighter in the condensed than in the dilute phase (**Figures 1** Error! Reference source not found.**E** and **S4A**).

Next, to assess which parts of Bik1 are required for the protein to undergo phase separation, we prepared two variants of Bik1 containing truncations in its N- or C-terminal region (**Figure 1 Error!** R**eference source not found.C**). Deletion of the CAP-Gly or the zinc-finger-containing domain (Bik1 ΔCG and Bik1 Δtail, respectively) both suppressed protein condensation at low salt conditions (e.g., 250 mM sodium chloride) where Bik1 FL readily formed droplets. Therefore, we investigated whether mixing the two truncated variants under low salt conditions can rescue their potential to undergo condensation. However, no visible droplets were observed under the same conditions used for Bik1 FL (**Figure S3**). This finding indicates that the introduced truncations in both Bik1 variants reduced the multivalency of the system needed for efficient phase separation of Bik1 FL. Additionally, we examined the relevance of the C-terminal EEY/F-like motif QQFF of Bik1 for the protein’s ability to undergo condensation. Interestingly, the deletion of the motif in Bik1 FL (Bik1 ΔQQFF; **Figure 1C**) was sufficient to prevent phase separation of this minimally truncated Bik1 variant (**Figure S3**).

Together, these results suggest that Bik1 is an elongated, complex multidomain homodimer comprising several folded domains and flexible regions. They further indicate that multiple intermolecular interactions involving Bik1’s CAP-Gly and tail domains are important for its phase separation. Notably, the possible multivalent interactions mediated by these two Bik1 elements with each other could establish “bridges” between several molecules, leading to the formation of branched, network-like structures (**Figure S4B**).

### Size-exclusion chromatography followed by small angle X-ray scattering analysis of Bik1

To experimentally assess the conformation of Bik1 FL in solution, we performed size-exclusion chromatography followed by small-angle X-ray scattering (SEC-SAXS) experiments.^30^ In addition, we analyzed Bik1 ΔQQFF, which is not able to undergo phase separation, and the coiled-coil domain of Bik1 (Bik1 CC; **Figure 1C**) to ascertain its contribution to the overall conformation of Bik1 FL and Bik1 ΔQQFF.

Examples of scattering profiles obtained in the high-salt buffer (500 mM sodium chloride) recorded for all three variants are shown in **Figure 2 Error! Reference source not found.A**. The pair distance distribution function, P(r), revealed similar values of radius of gyration, Rg, of 90-98 Å for Bik1 FL, 94-97 Å for Bik1 ΔQQFF, and 79-83 Å for Bik1 CC, as well as similar values of the maximal particle dimension, Dmax, for all variants of 280-320 Å, 294-320 Å, and 250-298 Å, respectively (**Figure 2 Error! Reference source not found.B, Table S1**). Notably, the Dmax values are consistent with the predicted length of the fully elongated Bik1 coiled-coil domain (∼300 Å; **Figure 1A**). Kratky analysis of the data (**Figure S5A**) suggests that the measured proteins are elongated and may include some unstructured elements.

**Figure 2.**
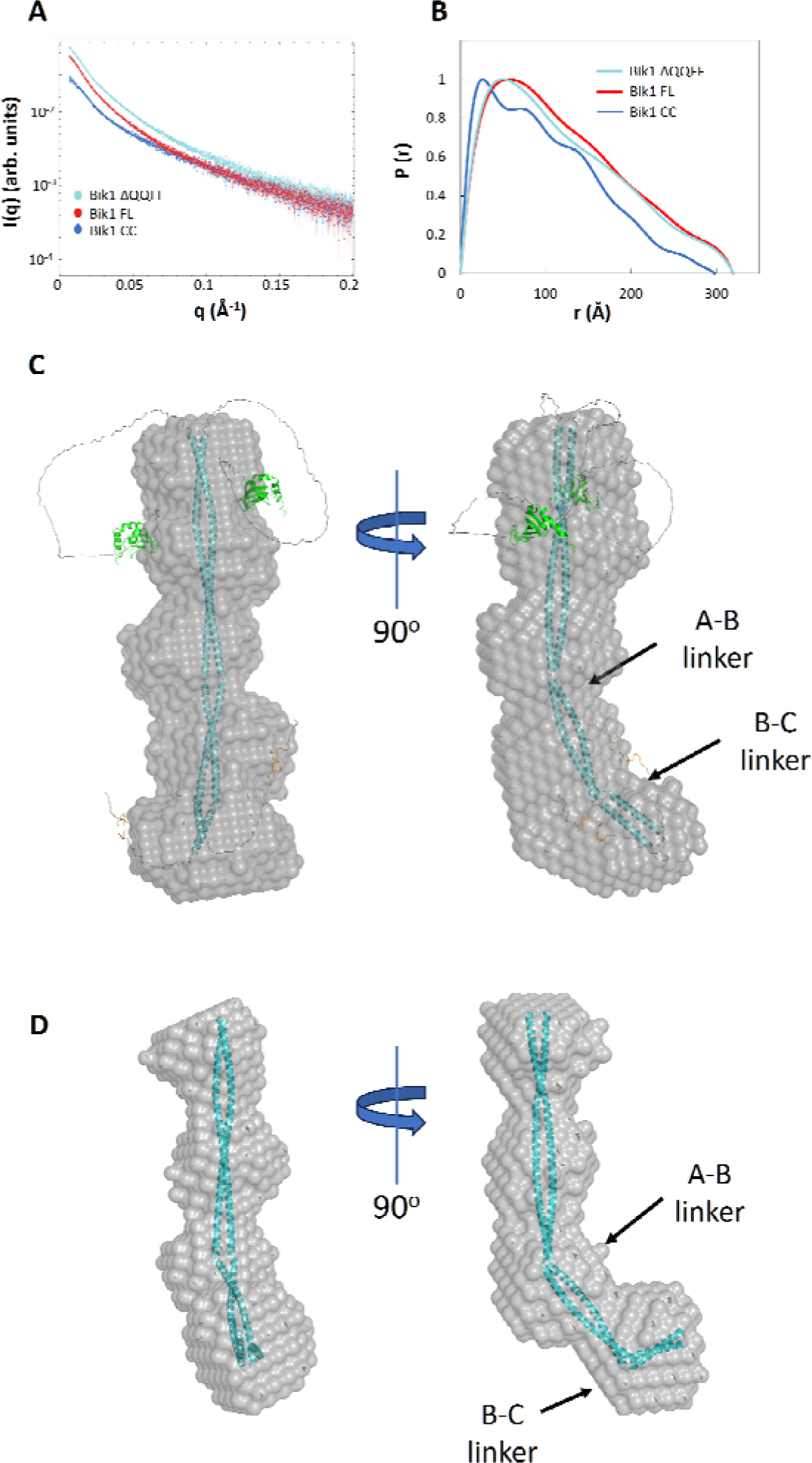
SEC-SAXS analysis of Bik1 variants. A) Representative scattering profiles obtained for Bik1 FL, Bik1 ΔQQFF, and Bik1 CC in a high salt buffer (20 mM Tris-HCl, pH 7.5, supplemented with 500 mM NaCl, 2% glycerol, and 1 mM DTT). B) Representative distance distribution functions obtained for Bik1 FL, Bik1 ΔQQFF, and Bik1 CC. C) A DAMMIF-generated molecular envelope (beads model) of Bik1 FL superimposed with its AlphaFold predicted model shown in Figure 1A and adjusted to fit the beads model. D) The molecular envelope of Bik1 CC superimposed with the Bik1 AlphaFold model truncated to the coiled coil and adjusted to fit the beads model. Two kinks suggested by the shape of the calculated envelope are marked with black arrows. See also **Figure S5**.

The bead-based envelope of the Bik1 FL dimer at a resolution of 61 ± 4 Å was calculated *ab initio* by averaging the 19 most similar models (out of 20 generated), resulting in a normalized spatial discrepancy (NSD) value of 0.72 ± 0.06, indicating a level of convergence in the model calculations (**Figure 2 Error! Reference source not found.C**). Similar envelopes were calculated for Bik1 ΔQQFF (resolution 65 ± 5 Å; NSD value of 0.71 ± 0.05) and Bik1 CC (resolution 43 ± 3; NSD value of 0.61 ± 0.04) by averaging 19 models (out of 20 generated) for each variant (**Figure S5BC**). The molecular envelope of Bik1 FL indicates that the Bik1 dimer adopts an elongated conformation with presumably two distinctive kinks, one in the middle part of the envelope and another one at its extremity. These two kinks could potentially be attributed to the predicted flexible linkers between coiled-coil segments (residues 298-302 and 358-362; **Figures 1A** and **S1**). Notably, the envelope obtained for Bik1 CC also displays two kinks at the same positions (**Figures 2D** and **S5**), implying that the coiled-coil domain of Bik1 predominantly influences the overall elongated shape of Bik1 FL’s envelope. The connection between the CAP-Gly domain and the coiled coil is mediated by the unstructured linker that holds the potential for diverse spatial arrangements. Consequently, akin to the C-terminal tail, these domains do not significantly contribute to the averaged overall shape of the calculated envelope. Worth noting is that the SEC-SAXS-generated molecular envelopes do not provide support for the “V-shaped”, folded-back conformations seen in certain predicted Bik1 FL structures (**Figure S1**).

Together, these findings are consistent with full-length Bik1 being a predominantly elongated dimer in solution, whose overall conformation is dominated by its two-stranded, parallel coiled-coil domain. They further indicate that both the N-terminal CAP-Gly and C-terminal zinc-finger containing domains are connected to the coiled coil with flexible linkers.

### Standard chemical crosslinking analysis of Bik1 in its dilute and condensed states

To investigate Bik1 interactions in its dilute and condensed states, we employed chemical crosslinking followed by mass spectrometry to identify crosslinked peptides.^31,32^ In our experiments, we used two crosslinking reagents, disuccinimidyl suberate (DSS) and pimelic acid dihydrazide (PDH).^33,34^ DSS reacts with primary amines (side chains of lysines, N-termini of proteins), which are separated by approximately 10-30 Å (Cα-Cα distance); PDH reacts with acidic amino acids (side chains of aspartates and glutamates), but can also form zero-length crosslinks in which amino and carboxyl groups are directly linked (Cα-Cα distance of approximately 7-30 Å). These reactions can be used to probe protein structures by introducing spatial restraints akin to a molecular ruler determined by the choice of the crosslinking reagents.^35^ During the crosslinking experiments, we identified the following four types of crosslinked peptides, which we define as follows:

i. Crosslinks: when a covalent bond is formed between two different residues, providing evidence of their proximity.
ii. Monolinks: when an amino acid forms a covalent bond with a hydrolyzed molecule of the crosslinking reagent, serving as a proxy for residue flexibility.
iii. Selflinks: when two amino acids at the same position form a covalent bond, indicating the presence of oligomers.
iv. Zerolinks: when two amino acids are covalently bonded without the presence of a spacer arms (linker).

Using the DSS reagent, we performed crosslinking reactions in three replicates with dilute single-phase (500 mM sodium chloride; referred to as “dilute” Bik1 from here onwards) and phase-separated (125 mM sodium chloride) Bik1 FL samples to study topological differences in interactions between Bik1 domains in both states. Since Bik1 FL forms stable homodimers in solution, we were not able to distinguish between intra- and intermolecular crosslinks, except for intra-dimer selflinks. Employing a conservative filtering strategy based on spectra quality (ld.Score > 25 and FDR < 0.05) and consistency (crosslinked peptides must be identified in at least two out of three replicates), we identified 70 unique crosslinked peptides for phase-separated Bik1 FL (low salt condition; 44 crosslinks, 21 monolinks, and 5 selflinks; **Figure 3A**; **Table S2**) and 60 unique crosslinked peptides for dilute Bik1 FL (high salt condition; 36 crosslinks, 22 monolinks, and 2 selflinks; **Figure 3B, Table S2**). To account for potential artifacts, different crosslinking reagent concentrations and distinct fragmentation conditions were tested (see **Materials and Methods**).

**Figure 3.**
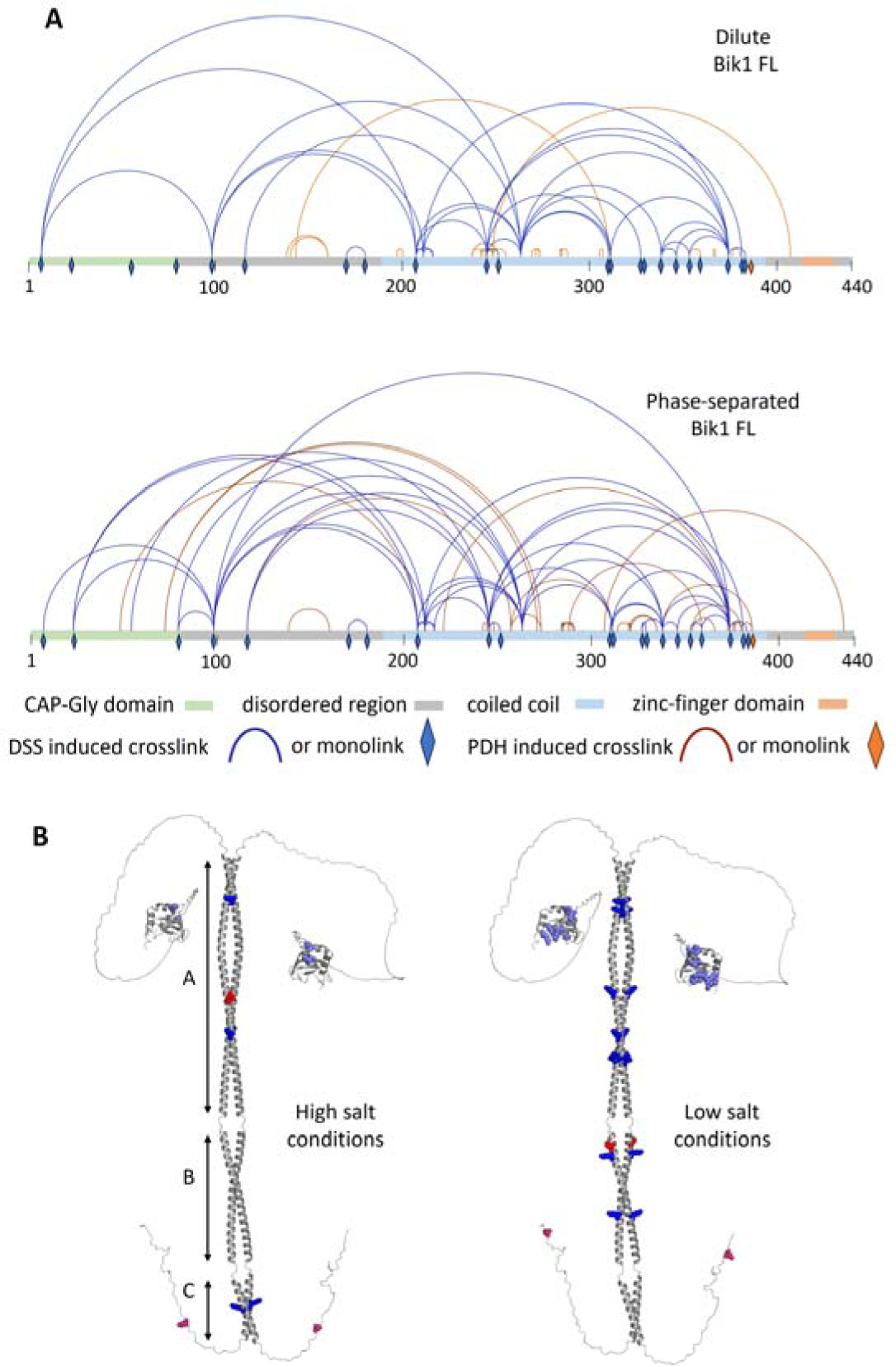
Inter- and intra-domain interactions identified in dilute and phase-separated Bik1 FL samples using XL-MS. A) Crosslinks and monolinks identified in dilute (10 mM HEPES, pH 7.5, supplemented with 500 mM NaCl) and phase-separated (5 mM HEPES, pH 7.5, supplemented with 250 mM NaCl) Bik1 FL samples. Crosslinks linking different regions of the protein are depicted as blue (DSS) or orange (PDH) lines; monolinks are shown as blue (DSS) or orange (PDH) rhombi. Bik1 domains are color-coded as follows: CAP-Gly domain in light green; disordered region in gray; coiled coil in light blue; zinc-finger domain in beige. B) Several interactions between the CAP-Gly domain (residues marked in light blue) and different regions of the coiled coil (residues marked in dark blue), or the C-terminal tail region (residues marked in pink) and different regions of the coiled coil (residues marked in bright red) have been identified in high and low salt conditions using both reagents, DSS and PDH. Interactions of the coiled coil with flanking domains cover crosslinks identified in dilute Bik1 FL, supernatant Bik1 FL, and Bik1 ΔQQFF (high salt conditions), or in phase-separated Bik1 FL and pellet Bik1 FL (low salt conditions; **Figure S6**). Note that due to the homodimeric nature of Bik1, inter- and intradimer interactions cannot be distinguished.

Differential analysis of mass spectra revealed an increased number of crosslinks and selflinks in phase-separated Bik1 samples, while the opposite trend was found for monolinks (**Figure S8C**). As the abundance of monolinks is proportional to the flexibility/exposition of the residue to the solvent,^36^ these results suggest a higher propensity of Bik1 to form a multivalent interaction network at low salt conditions. Four out of five selflink peptides were identified in the coiled-coil domain (Lys207, Lys245, Lys263, and Lys374), indicating the involvement of this region in protein dimerization. Moreover, we found multiple crosslinks between nearby residues of the coiled coil, which additionally support our computationally predicted Bik1 structure showing a parallel, in-register organization of the two-stranded coiled-coil domain (**Figure 1A**). Furthermore, based on the analyzed crosslinks (Lys7-Lys207, Lys7-Lys263, Lys23-Lys207, Lys23-Lys211, Lys54-Lys245, and Lys79-Lys263), we concluded that the CAP-Gly domain could interact with different regions of the coiled coil within or between Bik1 dimers (**Figure 3AB**). Interestingly, only two interactions involving the CAP-Gly domain and the coiled coil (Lys7-Lys207 and Lys7-Lys263) are specific for the dilute protein, while all other interactions are specific for phase-separated samples. These are most likely weak, nonspecific electrostatic interactions between the predominantly positively charged CAP-Gly domain and the negatively charged patches of the coiled coil (**Figure 1B**). The increased count of crosslinks observed between the CAP-Gly domain and the coiled-coil domain in the condensed phase (4 versus 2), coupled with the finding of a solution-specific monolink of Lys55, indicates a spatial rearrangement of the CAP-Gly domain that is accompanied by a reduction of its flexibility within phase-separated Bik1 samples.

Crosslinks identified using DSS chemistry cover most of the Bik1 sequence, excluding its C-terminal region containing the zinc-finger domain (residues 390-440). This region is marked by high flexibility, numerous acidic residues, and a lack of lysine and arginine residues that are targeted by DSS. To address this limitation, we used the PDH crosslinking reagent in combination with different proteases (AspN, GluC, chymotrypsin, and a combination of AspN and GluC). Utilizing this approach, we identified 59 unique crosslinked peptides with the ld.Score > 25 and FDR < 0.05 threshold, comprising 47 crosslinks, 2 monolinks, and 10 zerolinks. Although the efficiency in the identification of crosslinks with PDH was significantly lower than in the case of DSS, this orthogonal approach allowed us to study interactions between the C-terminal zinc-finger-containing domain of Bik1 and different regions of the protein’s coiled coil. For instance, we detected crosslinks between residues Asp407 and Asp248 for dilute Bik1 FL (**Figures 3AB** and **S8D; Table S3**). The distance between these residues measured using the AlphaFold model presented in **Figure 1A** is approximately 220 Å, which is much more than the typical distance between residues connected with this crosslinker (PDH; 7-30 Å).^34^ This event can be explained only by a significant back-folding of the Bik1 dimer or intermolecular interactions. In the phase-separated samples, we observed crosslinks from similar regions involving residues Asp435 and Glu307. These interactions would require kinks in the lower part of the coiled coil, as suggested by our computational predictions (**Figures 1A** and **S1**) and the SEC-SAXS-derived envelope (**Figure 2C**), and then the flexible C-terminal tail region could reach Glu307. Also, additional interactions between the CAP-Gly domain and the coiled coil were observed in phase-separated samples: Asp48-Lys207, Lys72-Glu271, and Lys72-Glu273 (**Figure 3B**). Remarkably, coiled-coil residues that were observed to interact with the CAP-Gly domain or with the C-terminal tail region are located close to each other (e.g., Asp248 and Lys245). Therefore, this analysis suggests that both the N-terminal CAP-Gly domain and the C-terminal tail region of Bik1 can come close to each other to interact. However, direct interactions between the C-terminal tail and the CAP-Gly domain have not been determined in our crosslinking experiments.

Together, these results illustrate the great multivalent nature of the multidomain Bik1 dimer and suggest a higher propensity for multivalent interactions in phase-separated Bik1 samples. Moreover, the identified interactions between the CAP-Gly and coiled-coil domains indicate spatial rearrangements of the protein occurring during its condensation.

### Quantitative chemical crosslinking analysis of phase-separated Bik1

To assess the dynamics inside phase-separated Bik1 samples, we adopted a quantitative approach to measure the intensity of crosslinked peptides. This approach enabled us to study changes in residue distances and the dynamical reorganization of the protein during condensation. A similar strategy has previously been successfully used to characterize conformational changes of proteins in various contexts, such as the characterization of the ubiquitin ligase E6AP complex,^37^ investigations of single site mutation effects on a protein structure,^38^ and studies of FUS dynamics in the context of phase separation.^35^ For our experiments, we devised a two-step approach: i) identification of crosslinked peptides that characterize specific structural states (denoted “conformospecific” crosslinks), and ii) quantification of crosslinked peptides via targeted proteomics and differential analysis (**Figure 4A**). For the quantitative analysis, we harnessed the sensitivity and the high quantification accuracy associated with intensity-based fragment analysis of a targeted acquisition scheme (Parallel Reaction Monitoring, P.R.M.).^39,40^

**Figure 4.**
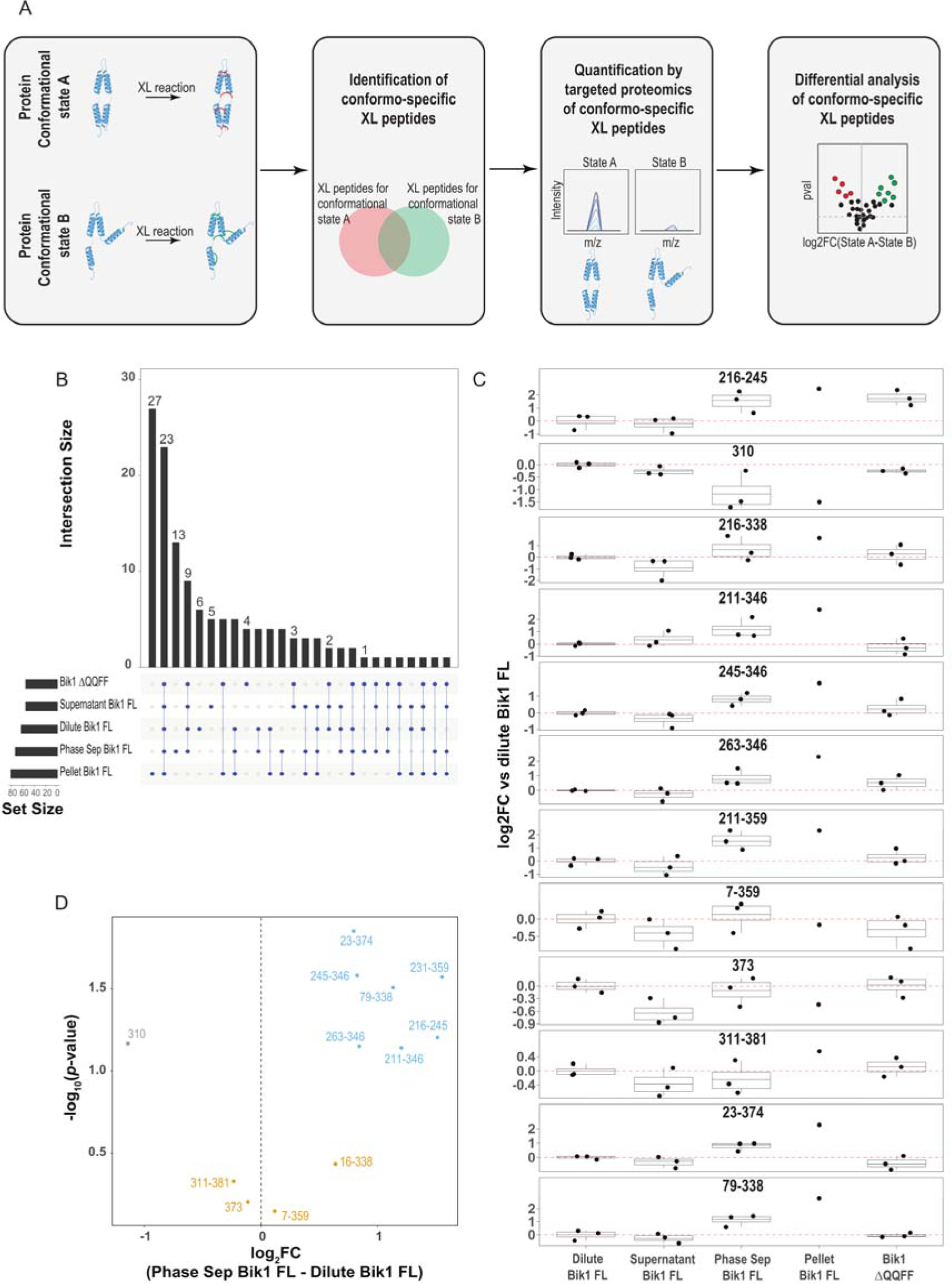
Abundance of crosslinks across different Bik1 samples. A) Quantitative workflow to probe Bik1 conformational changes. Crosslinked (XL) peptides were obtained from Bik1 FL and Bik1 ΔQQFF: dilute Bik1 FL (500 mM NaCl), Bik1 ΔQQFF (500 mM NaCl), phase-separated Bik1 FL (125 mM NaCl), and phase-separated Bik1 FL centrifuged before the reaction with DSS and separated into supernatant and pellet. Crosslinks specific to one condition (conformospecific crosslinks) are then employed to monitor structural rearrangements in Bik1 samples using quantitative targeted proteomics. B) Upset plot of identified crosslinks across different conditions (dilute Bik1 FL, supernatant, phase-separated Bik1 FL, pellet, Bik1 ΔQQFF). C) Boxplot showing the abundance of crosslinked peptides normalized for Bik1 FL abundance at different conditions (pellet, dilute Bik1 FL, supernatant, Bik1 ΔQQFF, phase-separated Bik1 FL). Each dot represents the results for a replicate (N=3 for all conditions, except for pellet where N=1), with the box plot boundaries indicating the quantiles Q1 (25%) and Q3 (75%), and the average value denoted by a line across the box. D) Differential abundance levels for conformospecific crosslinks in dilute and phase-separated samples. The volcano plot shows the log2 fold changes of crosslinks abundance and their corresponding statistical significance (two-sided unpaired Student’s t-test, N=3 technical replicates) between the two conditions.

To identify conformospecific crosslinks of Bik1, we used DSS as a crosslinking reagent and trypsin for proteolysis. We probed the following states: Bik1 FL at 500 mM NaCl (dilute protein), Bik1 FL at 125 mM NaCl (phase-separated protein), and Bik1 ΔQQFF (at 500 mM NaCl) that cannot undergo phase separation. Additionally, we prepared a series of Bik1 FL phase-separated samples that were centrifuged, and the reaction with DSS was performed separately for the condensed (pellet) and dilute (supernatant) phases. Using the same crosslinking filtering criteria as described above, we identified a total of 199 crosslinked peptides, including 159 crosslinks, 29 monolinks, and 11 selflinks (**Figure S6**; **Table S2**). As shown in **Figure 4B**, we observed a significant overlap in identified peptides across most conditions, except for the Bik1 FL pellet. From the identified crosslinks, we selected for the targeted analysis only those that are characteristic for specific conditions (conformospecific crosslinks) for the targeted analysis. For instance, Lys211-Lys346, Lys211-Lys359, Lys216-Lys245, Lys216-Lys338, Lys79-Lys338, and Lys263-Lys346 were identified only in the Bik1 FL pellet condition, while Lys245-Lys346 and Lys23-Lys374 are specific to the BIK1 ΔQQFF sample and the Bik1 FL supernatant condition.

For the quantitative analysis, we tracked 12 crosslinked peptides, normalizing their intensity for the abundance of Bik1 protein to account for potential variations of protein levels. For the normalization, we used the two not crosslinked “householder” peptides MVLEEVQPTFDR and HSGNQQSMDQEASDHHQQQEFGYDNR (**Figure S7D**). The quality control for the targeted analysis experiment is shown in **Figure S7ABC**. Unsupervised hierarchical clustering of crosslinked peptide intensities revealed that pellet and phase-separated conditions differed significantly from the other conditions, forming a separate cluster (**Figure S7E**). This result indicates that these two conditions share the same residue interaction network.

The identified crosslinks exhibit different abundance levels and can be divided into the following three groups (**Figure 4C**; **Table S4**):

I. Crosslinks exhibiting higher intensity in the pellet and phase-separated samples than in the supernatant and dilute conditions (e.g., Lys216-Lys245, Lys211-Lys346, Lys245-Lys346, Lys263-Lys346, Lys211-Lys359, Lys23-Lys374, and Lys79-Lys338). Interestingly, the abundance levels of these crosslinks in the Bik1 ΔQQFF samples are between the levels recorded for the dilute and phase-separated samples, which might suggest an intermediate behavior with the formation of higher oligomers in the solution but not condensation leading to the formation of micrometer-sized phase-separated droplets.
II. Monolinks with reduced abundance levels in the pellet sample (Lys310 and Lys373). They can serve as proxies for assessing residue accessibility. This result suggests that in the Bik1 pellet condition, the coiled-coil region around Lys310 and Lys373 is much less exposed to the solvent than in other conditions. Differences in the coiled-coil residue exposure to the solvent or other Bik1 domains might be related to the presence of the kinks between coiled-coil segments, as computationally predicted and revealed by the SEC-SAXS-derived envelopes (**Figures 1A** and **2D**).
III. The crosslinked peptides Lys216-Lys338, Lys7-Lys359, and Lys311-Lys381 maintain constant abundance levels across the phase-separated, dilute, supernatant, and Bik1 ΔQQFF samples (see above for exact sample designation).

The observed differences in the abundance of the identified crosslinked peptides can be used to characterize the most significant conformational changes of Bik1 FL during the protein’s transition between its dilute and phase-separated states. The Volcano plot (**Figure 4D**) revealed that inter- and intramolecular interactions within the coiled-coil domain (Lys359-Lys231, Lys245-Lys347, Lys231-Lys359, Lys216-Lys245, Lys263-Lys346, and Lys211-Lys346) and with the CAP-Gly domain (Lys23-Lys374 and Lys79-Lys338) are between 1 and 1.5 log2 fold (approximately 1.5 and 2.9 times) more abundant in the low salt (phase-separated protein) than in the high salt (dilute protein) condition. On the contrary, the monolink of Lys310 exhibits a 1 log2 fold higher level (approximately 2.2 times greater abundance) in the dilute state, suggesting higher exposure of this coiled-coil domain to the solvent.

Together, these results provide insight into the conformational rearrangements of Bik1 dimers occurring during their transition from the dilute to the phase-separated states. They further underscore the essential contribution of all three coiled-coil segments as well as their interactions with the CAP-Gly domain to create an intricate multivalent network of interactions.

### Small-angle X-ray scattering analysis of phase-separated Bik1

Next, we investigated the supramolecular structure of Bik1 condensates. To this end, SAXS measurements were conducted on phase-separated Bik1 FL droplets as they flowed through a microfluidic chip. Bik1 droplets showed enhanced scattering at low q (large length scales), which was absent from the dilute phase scattering (**Figure 5A**). In this regime, the SAXS signal decays as I(q) Ill q^−2^, suggesting the presence of a fractal network with a fractal dimension *d* ≈ 2. This fractal structure extends from the lowest *q* reached in the SAXS data (from > 3000 Å) down to about 300 Å (**Figures 5A** and **S9A**). To visualize this supramolecular structure, we generated multiple real-space structures that were consistent with the scattering data. In all cases, thousands of copies of a simplified Bik1 structural model were placed in a spherical volume with a diameter of 1 µm (see **Materials and Methods** for a detailed description of the approach). The positions and orientations of these model copies were iteratively repositioned to fit the SAXS data in the range 0.002 Å^−1^ < *q* < 0.03 Å^−1^ corresponding to lengths starting from about 200 Å up to 3100 Å (**Figures 5A**). We considered a range of Bik1 models, including a Gaussian blob and three different spherocylinder models that approximate the SEC-SAXS solution structure of the protein (**Figures 5A** and **S9B**). In all cases, we achieved a good agreement with the scattering data and example structures, as shown in **Figure 5C, Figure S9C,** and **Movie 1**. As expected for a fractal network, we observed a heterogeneous distribution with protein-rich regions separated by protein-free (solvent-rich) voids.

**Figure 5.**
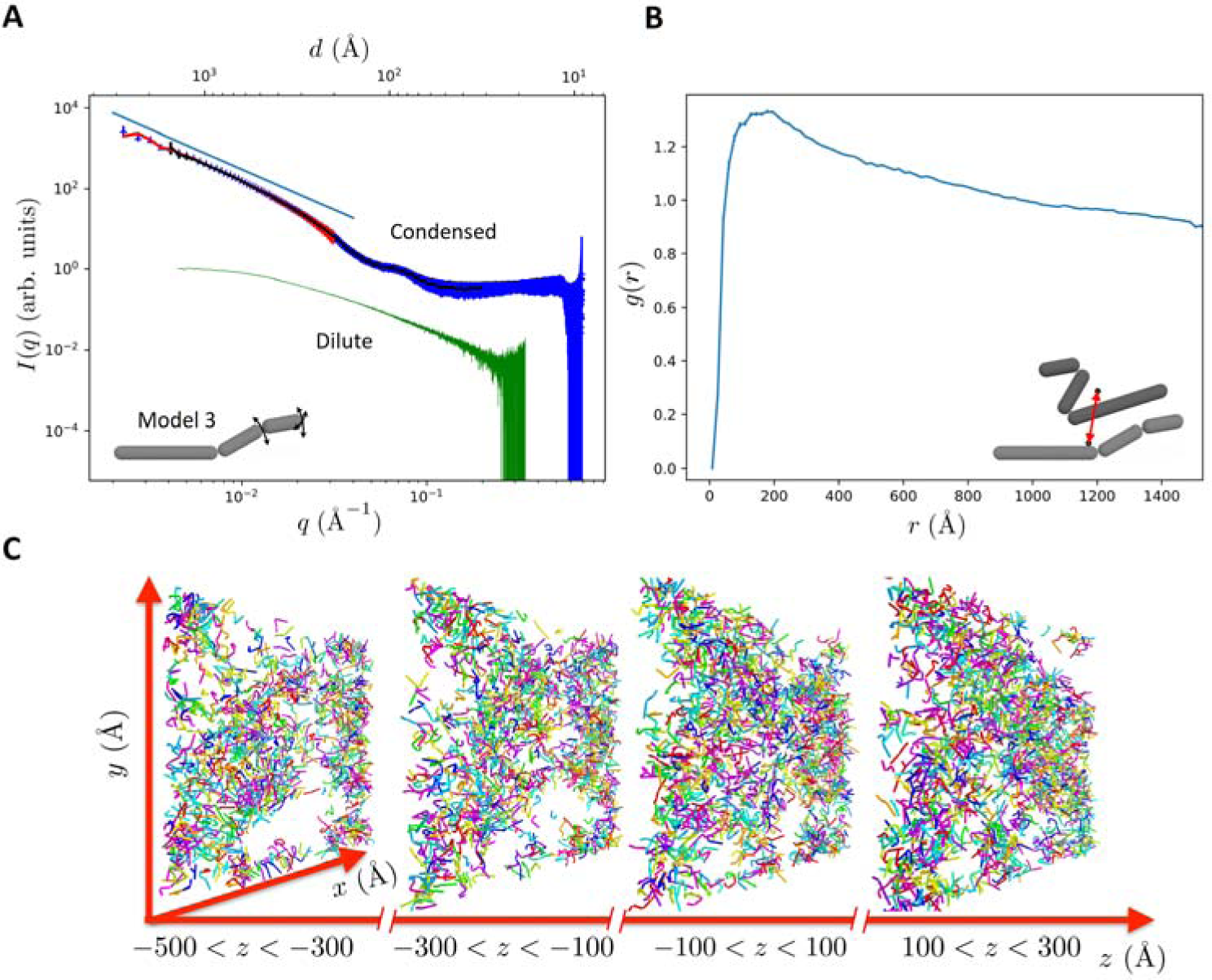
SAXS analysis of phase-separated Bik1. A) SAXS measurement (blue curve) with Gaussian-blob fit (red curve) and fit obtained with 50’000 model 3 copies (black curve; see **Materials and Methods** for model description and fitting procedure). The model 3 fit is restricted to q > 2π/σ_w_ due to the Gaussian window with σ_w_ = 1500 Å; it was used to model the fractal structure. The straight blue line above the SAXS measurement represents the *I(q)* ∼ *q^−2^* behavior expected for a fractal network with a fractal dimension of 2. The green curve shows the Bik1 form factor obtained from our SEC-SAXS data (Figure 2A). Inset, schematic representation of model 3. B) Radial distribution function of the centers of 50’000 model 3 copies. The decay for *r* larger than the nearest-neighbor peak is expected for a fractal network. Inset, two model 3 copies with their centers of mass (dots). The arrow highlights the distance *r* between their centers. C) Sections taken from the fractal structure obtained with 50’000 model 3 copies.

To test for translational order, we calculated the radial distribution function, *g*(*r*), from the centers of each Bik1 model in the fitted structure (**Figure 5B**). The peak near 200 Å reflects the typical distance between nearest neighbors and the decay at larger distances continues smoothly to a plateau *g*(*r)*=1, which is expected for a fractal network. To test for orientational order, we calculated the nematic order parameter of the fitted supramolecular structure. The nematic order parameter is a local measure of the relative orientation - it is zero in the absence of orientational order when all molecules have random orientations and one when all molecules are aligned in the same direction. We calculated the nematic order parameter in cubic boxes with side lengths of 400 Å. Since the number of particles in the box fluctuates due to their heterogeneous distribution, we used a lower cutoff of 30 particles in the box to obtain a reliable order parameter. In these boxes, we find the order parameter to be below 0.4 in all cases (**Figure S10**). This suggests that Bik1 molecules tend to align but nevertheless remain largely in an isotropic state. With an aspect ratio of 12.5, Bik1 molecules could organize into a strongly aligned (liquid crystalline) state at volume fractions above 0.18.^41^ This is about a factor of two above the experimentally estimated volume fraction of Bik1 in condensates. ^20^

Together, these results indicate that phase-separated Bik1 dimers form a fractal network, characterized by protein-dense and protein-free regions. They further indicate an isotropic distribution of Bik1 dimers in the condensed phase.

## Discussion

The ubiquitous nature of protein liquid-liquid phase separation has come to the forefront in recent studies, elucidating its fundamental involvement in critical cellular processes.^14–17,42^ Compelling evidence is now surfacing, shedding light on the integral role of phase separation in shaping human health and its involvement in the onset of diseases.^12,43^ However, the molecular mechanisms of liquid-liquid phase separation and structures of condensed proteins remain poorly understood, particularly when it comes to multidomain proteins. Here, we structurally characterized Bik1 in its dilute single-phase and phase-separated states.

To better understand which domains of Bik1 are essential for the protein to undergo condensation, we produced several variants containing truncations in N- and C-terminal parts of the protein. Our biochemical studies revealed that the CAP-Gly domain and the C-terminal tail are critical for the condensation of Bik1. The smallest change to the protein’s sequence that results in the loss of phase-separation properties was the deletion of the C-terminal EEY/F-like motif QQFF. The CAP-Gly domain of Bik1 was previously shown to mediate its interactions with α-tubulin, Bim1, and itself.^44^ Based on our results, we hypothesize that under reduced ionic strength conditions, the interaction between the N-terminal CAP-Gly domain and the C-terminal tail might bridge several molecules, leading to the formation of branched, network-like structures.

A combination of computational and SAXS methods allowed us to obtain structural information regarding the overall conformation of the Bik1 homodimer in solution. In the past, SAXS has been demonstrated as a robust and effective method for investigating protein phase separation.^30^ Our results suggest that Bik1 adopts a predominantly elongated conformation with two kinks caused by discontinuities in the heptad-repeat sequence of its two-stranded, parallel coiled-coil domain. The SAXS-derived beads models obtained for Bik1 FL and Bik1 CC are very similar to each other and, therefore, show that the overall shape of the Bik1 dimer is dominated by the elongated conformation of the coiled-coil domain. However, we cannot exclude the possibility that some Bik1 dimers adopt a backfolded conformation. The N-terminal CAP-Gly and C-terminal zinc-finger containing domains were predicted to be connected to the coiled coil via flexible linkers and, therefore, have the flexibility to interact with different coiled-coil segments, especially patches with opposite charges. Using chemical crosslinking followed by mass spectrometry, we confirmed multiple interactions of all three coiled-coil segments with both the CAP-Gly domain and the C-terminal flexible tail. Since N- and C-terminal domains engage with comparable regions of the coiled coil, they also have the potential to interact with each other. In this context, Bik1’s mammalian orthologue, CLIP-170, can fold back upon itself due to interactions between its N- and C-terminal domains.^19,45^ Notably, CLIP-170 was also recently reported to form liquid condensates *in vitro* and in cells.^45^ It was hypothesized that phase-separation of CLIP-170 facilitates the concentration of several protein partners, including tubulin, at the ends of microtubules, an activity that contributes to the regulation of microtubule dynamics in cells.^45^

The crosslinks identified for the phase-separated Bik1 FL underscore the essential contribution of all three coiled-coil segments, as well as their interactions with the CAP-Gly domain, creating an intricate multivalent network of interactions. Furthermore, differences in crosslinks observed in low and high salt conditions indicate spatial rearrangements of the protein during condensation. To study these spatial rearrangements, we designed a quantitative crosslinking approach, which revealed that specific interactions within coiled-coil segments or with the CAP-Gly domain (e.g., Lys23-Lys374) are between 1.5 and 2.9 times more abundant in the condensed state of the protein. We also identified monolinks with reduced abundance in low salt conditions (e.g., Lys310 and Lys373), suggesting that these regions are also involved in interactions during protein phase separation. Interestingly, the abundance levels of the tracked crosslinks in the Bik1 ΔQQFF samples fall within a spectrum between those observed in the dilute and phase-separated Bik1 FL samples. This might suggest the formation of higher oligomers in solution; however, without progressing to the extent of condensation that would result in the formation of micrometer-sized, phase-separated droplets.

Finally, we investigated the internal structure of phase-separated Bik1 droplets. Using confocal fluorescence microscopy, we observed that the fluorescently labeled Bik1 molecules are homogenously dispersed at the micron-scale. However, our SAXS data suggests that Bik1 droplets are heterogeneous at the supramolecular scale, from 30 to 300 nm. At this level of organization, Bik1 condensates form a network with a fractal dimension of about 2. Notably, fractal networks are a hallmark of percolation processes. In percolation, a system spanning network is formed through the binding of building blocks. Percolation models have a long history in the study of polymer sol-gel transitions,^46^ and more recently, they have been proposed to play an essential role in biomolecular condensates.^47,48^ Our observation of a fractal network within Bik1 droplets provides direct structural evidence for percolation in fluid biomolecular condensates. Based on these findings, we hypothesize that the fractal structure enables Bik1 dimers to achieve a high viscosity while maintaining a relatively low protein concentration in their condensed phase.^20^

In summary, our study provides a deeper understanding of structural changes and mechanisms involved in the phase separation of a complex multidomain, oligomeric protein. We present an experimental framework for investigating the supramolecular structural organization of phase-separated protein networks. In our model system, we found structural evidence supporting recent models emphasizing the role of percolation in biomolecular condensates. Gaining insight into how proteins arrange themselves within complex condensed phases will significantly enhance our understanding of the role of protein phase separation in fundamental cellular processes.

## Materials and Methods

### Protein purification

The DNA encoding *S. cerevisiae* Bik1 (Uniprot ID: P11709) and its variants (mNG-Bik1 FL, Bik1 FL, Bik1 ΔCG, Bik1 Δtail, Bik1 ΔQQFF, Bik1 CC, Bik1 CG) were cloned into the pET-based bacterial expression vector PSPCm2, which encodes for the N-terminal 6xHis-tag followed by a PreScission cleavage site using a positive selection method.^49^

All protein production was performed in the *E. coli* strain BL21-CodonPlus (DE3)-RIPL in an LB medium containing 50 μg/ml of kanamycin. When the cultures reached an OD_600_ of 0.6 at 37°C, they were cooled down to 20°C, induced with 0.75 mM isopropyl 1-thio-β-D-galactopyranoside (IPTG), and shaken for another 16 hours at 20°C. After harvesting by centrifugation, the cells were sonicated in the presence of protease inhibitors (cOmplete cocktail; Roche) and 0.1% bovine deoxyribonuclease I in purification buffer (20 mM Tris-HCl, pH 7.4, supplemented with 500 mM NaCl, 10 mM imidazole, and 2 mM β-mercaptoethanol).

Proteins were purified by immobilized metal-affinity chromatography (IMAC) on a HisTrap HP nickel-Sepharose column (GE Healthcare) at 4°C following the manufacturer’s instructions. The column was equilibrated in IMAC buffer A (20 mM Tris-HCl, pH 7.4, supplemented with 500 mM NaCl, 10 mM imidazole, and 2 mM β-mercaptoethanol). Proteins were eluted by IMAC buffer B (IMAC buffer A containing 400 mM imidazole in total). In the case of the Bik1 FL cleaved sample, the N-terminal 6xHis-tag was cleaved off by an in-house produced HRV 3C protease^50^ in IMAC buffer A for 16 hours at 4°C. Then, samples were reapplied on the IMAC column to separate the 6xHis-tag from the uncleaved protein.

Protein samples were concentrated and loaded onto a size exclusion chromatography (SEC) HiLoad Superdex 200 16/60 (mNG-Bik1 FL, Bik1 FL, Bik1 ΔCG, Bik1 Δtail, Bik1 ΔQQFF,) or Superdex 75 16/60 (Bik1 CC, Bik1 CG) columns (GE Healthcare), which were equilibrated in SEC buffer (20 mM Tris-HCl, pH 7.5, supplemented with 500 mM NaCl, 10% glycerol, and 1 mM DTT) or in XL buffer (10 mM HEPES, pH 7.5, supplemented with 500 mM NaCl and 1 mM DTT) for XL-MS experiments. In the case of Bik1 CG and Bik1 CC, the SEC buffer contained 150 mM NaCl final concentration. Bik1 protein-containing fractions were pooled and concentrated to the desired concentration. The protein quality and identity were assessed by SDS-PAGE and mass spectrometry, respectively.

### Circular dichroism (CD) spectroscopy

CD spectrum of Bik1 FL (0.2 mg/ml in 20 mM Tris-HCl, pH 7.4, supplemented with 500 mM NaCl) was recorded at 25°C on a Chirascan-Plus spectrophotometer (Applied Photophysics Ltd.) equipped with a computer-controlled Peltier element using a quartz cuvette of 1 mm optical path length. CD spectra were recorded between 200 and 260 nm in triplicates. Thermal unfolding profiles were recorded by CD at 222 nm by continuous heating at 1°C min^−1^.

### Fluorescence microscopy

10% mNG-Bik1 FL/90% Bik1 FL was diluted in buffer (20mM Tris-HCl, pH 7.4, supplemented with 250mM NaCl) to a final concentration of 3.4 mg/ml. The phase-separated droplets were deposited onto glass coverslips coated with PEG-Silane (Gelest) and sealed with a second coverslip with a spacer in between. Samples were imaged on an inverted confocal microscope (Nikon Ti2 Eclipse with Yokogawa CSU-W1 spinning disc) using a 60x water immersion objective (NA of 1.2).

For quantification of intensities inside droplets, acquired z-stacks were analyzed with a MATLAB code. Briefly, droplets were located above a threshold intensity in 3D, and the center z-plane for each drop was identified. In each mid-plane, the centroid of the droplet circle was identified, and intensities averaged azimuthally as a function of distance from the droplet surface. Radial intensities of 233 different droplets were then averaged to give a representative profile of mNeonGreen intensity as a function of distance from a droplet edge.

### Crosslinking mass spectrometry (XL-MS)

#### Crosslinking reactions

Bik1 FL and Bik1 ΔQQFF samples were subjected to a crosslinking reaction with 0.5 or 1 mM isotope labeled di-succimidylsuberate (DSS-d0 and DSS-d12; CreativeMolecules Inc.) following a previously described protocol.^33^ The reaction of Bik1 variants with the acid crosslinking reagent was initiated by the addition of a mixture consisting of pimelic acid dihydrazide (light and heavily labeled: PDH-d0, PDH-d10) at 8.9 mg/ml with DMTMM at 12mg/ml. The reaction was quenched by removing the reagents with a Zeba Spin Desalting column (0.5 mL, 7K MWCO, Pierce) following the previously published procedure.^34^ Upon quenching, samples were dried, dissolved in 8 M Urea, reduced with 5 mM TCEP, and alkylated with 10 mM iodoacetamide. The proteolysis was performed overnight at 37 ^O^C in 1M urea using different proteases (trypsin 1:50 E:S; AspN 1:200 E:S; AspN/GluC 1:100 E:S; GluC 1:100 E:S; chymotrypsin 1:100 E:S) and then quenched with 5% formic acid. Generated peptides were subjected to cleanup (C18 column, TheNest Group) and separated by SEC using an ÄKTA micro chromatography system (GE Healthcare) using a Superdex 200 Increase 3.2/30 column (Cytiva). The SEC fractions were then dried and re-dissolved in 5% acetonitrile and 0.1% formic acid for mass-spectrometry analysis. For the targeted analysis of crosslinked peptides, samples were not subjected to SEC fractionation.

#### Data acquisition

Liquid chromatography tandem mass spectrometry (LC-MS/MS) measurements were performed on an Orbitrap QExactive+ mass spectrometer (Thermo Fisher) coupled to an EASY-nLC-1000 liquid chromatography system (Thermo Fisher). Peptides were separated using a reverse phase column (75 µm ID x 400 mm New Objective, in-house packed with ReproSil Gold 120 C18, 1.9 µm, Dr. Maisch GmbH). Data acquisition was done in two modes: Data Dependent Acquisition (D.D.A.) and Parallel Reaction Monitoring (P.R.M.).

For the identification of crosslinking peptides in DDA mode, an LC method was set up to separate peptides across a 60 min gradient: from 5% to 25% in 55 min and from 25% to 40% in 5 min (buffer A: 0.1% (v/v) formic acid; buffer B: 0.1% (v/v) formic acid, 95% (v/v) acetonitrile). The acquisition method was performed with one MS1 scan followed by a maximum of 20 scans for the top 20 most intense peptides (TOP20) with MS1 scans (R=70’000 at 4001m/z, max IT= 64ms AGC=1e5), HCD fragmentation (NCE=25 or 28%), isolation windows (1.5 m/z) and MS2 scans (R=35’000 at 4001m/z, maxIT = 110ms, AGC= 5e4ms). A dynamic exclusion of 30 sec was applied, and charge states lower than three and higher than seven were rejected for the isolation.

For the targeted quantification of crosslinking peptides in P.R.M. mode, an LC method was set up to separate peptides across 95 minutes gradient: from 5% to 40% in 90 min and from 40% to 50% in 5 min (buffer A: 0.1% (v/v) formic acid; buffer B: 0.1% (v/v) formic acid, 95% (v/v) acetonitrile). The MS acquisition of targeted peptide (12 crosslinks heavy and light, 2 householder peptides for Bik1 peptides and 11iRT peptides) was set up with the combination of one MS1 untargeted scan (R=70’000 at 400m/z, maxIT = 100ms, AGC= 3e6) and 38 scheduled targeted scan (R=35’000 at 400m/z, maxIT = 110ms, AGC= 2e5) using an isolation window of 1.8m/z and HCD fragmentation (NCE=28%).

#### Peptide identification and data analysis

For the identification of crosslinking data, mass spectra were converted to mzXML format and searched with xQuest/xProphet^51^ against a database containing the FASTA sequence of Bik1 and its decoy sequence. Crosslinked peptides with ld.Score > 25, delta S > 0.95, and FDR < 0.05 were considered in the analysis. XL peptides were visualized using the xiVIEW web-based tool for the analysis of cross-linking results.^52^ Conformospecific crosslinked peptides (crosslinks identified only in specific conditions) were selected and analyzed using a targeted proteomics approach. The detailed procedure has been previously reported.^38,53^ We generated a library based on the previous xQuest identification; only common transitions from light and heavy crosslinked peptides were used for this analysis. The quantification of targeted peptides was manually performed using Skyline-daily^54^ based on the following criteria: (1) co-elution of heavy and light cross-linked peptides, (2) matching of the peak shape, and (3) matching intensity for at least six common transitions of heavy and light crosslinked peptides (transitions shared between the light and heavy form of the crosslinked peptide). The abundance of crosslinked peptides was then calculated by summing the integrated area of ten common transitions per peptide (five transitions for the heavy and light forms of the crosslinked peptide, respectively). Crosslinked peptides were normalized for the intensity of two non-crosslinked Bik1 peptides (MVLEEVQPTFDR and HSGNQQSMDQEASDHHQQQEFGYDNR). The significance of change for the log2 abundance of the crosslinked peptides was estimated with p values using a two-sided, not paired t-test.

To account for potential artifacts from crosslinking reagent concentration and detection based on different collision energy during fragmentation, we performed the crosslinking reaction of Bik1 FL in solution at different DSS concentrations (0.5mM and 1mM) and distinct fragmentation settings (NCE = 25 and 28). We identified 117 cross-linked peptides (90 intra-protein crosslinks and 27 monolinks) using a filtering cutoff of ld.Score > 20 and FDR< 0.05. We consistently identified 19 crosslinking peptides across all experimental conditions tested. Notably, we observed a higher degree of overlap in the results when conducting the crosslinking reaction at different concentrations while maintaining the same fragmentation energy, as compared to varying the fragmentation energy while keeping the concentration of the crosslinking reagent constant. As increasing the concentration of crosslinking reagents from 0.5 mM to 1 mM did not result in more self-crosslinked peptides, which are the intermolecular crosslinks that occur when a peptide forms a covalent bond with the same peptide, we proceeded to study differences in interactions between dilute and phase-separated Bik1 FL using 1mM DSS.

The entire dataset, including raw data, generated tables and scripts used for the data analysis are available in the PRIDE repository^55^: PXD050928: Effect of DSS concentration and fragmentation on the identification of XL peptides (**Table S5**); PXD050929: Identification of DSS XL peptides in Bik1 variants and different conditions (**Table S2**); PXD050930: Identification of PDH XL peptides in Bik1 at different conditions (**Table S3**); PXD050931: Quantification of DSS XL peptides in Bik1 variants and different conditions (**Table S4**).

### Size-exclusion chromatography followed by small-angle X-ray scattering (SEC-SAXS)

SEC-SAXS measurements were performed at beamline B21^56^ at the Diamond Light Source using the photon energy of 13.1 keV. A SEC step was performed for each sample prior to the SAXS measurement using a Shodex KW-404 column equilibrated in SAXS buffer (20 mM Tris-HCl, pH 7.5, supplemented with 500 mM NaCl, 2% glycerol, and 1 mM DTT) and connected in-line to the X-ray scattering measurement cell. Samples of Bik1 FL, Bik1 ΔQQFF, and Bik1 CC were injected at concentrations of 4, 6, and 8 mg/ml in 50 µl volumes and with a flow rate of 0.16 ml/min.

The ATSAS^57^ software was used to perform buffer subtraction, SEC peak scattering intensity summation, radius of gyration (Rg) calculation from Guinier plots, and analysis of distance distribution functions P(r). Ab initio calculation of molecular envelopes was performed using DAMMIF.^58^ Model averaging and pairwise cross-correlation were performed using DAMAVER.^59^ For all analyzed proteins, 20 bead models were calculated using random start seeds with no presumed internal symmetry (P1). For the final envelope, the most similar 19 models (out of 20) were averaged for each protein. The resolution of the calculated envelopes was estimated using SASRES.^60^ Graphic visualization of protein molecules was done using PyMOL (the PyMOL Molecular Graphics System, Version 2.4.1 Schrödinger, LLC).

### Small-angle X-ray scattering (SAXS) measurements of phase-separated Bik1 samples

#### Data collection

SAXS experiments were conducted at the cSAXS beamline at the Swiss Light Source (Paul Scherrer Institut). The monochromatic beam, with a photon energy of 11.2 keV, corresponding to a wavelength of 0.11 nm, was focused so that the full width at half maximum (FWHM) beam size at the sample position was 12×26 µm along and across the flow direction in the microfluidic channel (Fluidic 394, featuring a 200×200 μm channel, #10000016 Microfluidic ChipShop), respectively. This beam size was chosen to ensure the entire beam fits into the microfluidic channel without any edge scattering.

Scattered X-rays were detected using a Pilatus detector,^61^ positioned 2.172 m downstream of the sample, as determined by analyzing the scattering signal of a silver behenate standard. The flight path between the sample and detector was evacuated to minimize parasitic scattering and absorption. A beamstop served both to protect the pixel detector from the direct beam and to allow for measuring the intensity of the unscattered beam.

To mitigate radiation damage, pre-formed droplets of Bik1 FL (4 mg/ml) in 20 mM Tris-HCl, pH 7.4, buffer supplemented with 260 mM NaCl were introduced into a microfluidic chip at a pumping rate of 0.1 µl/s. To minimize sample consumption while maintaining high sample flow rates, the droplet solution was co-flowed with 2 x 0.1 µl/s of buffer, which were fed through the left and right channels in the microfluidic cross junction, leading to a sheet flow with a dilution of the sample along the channel. A few seconds after the channel was filled with the sample, data were acquired at 1.5 cm, 2 cm, and 4.5 cm from the mixing point, respectively. The corresponding signal from the buffer was measured at approximately the same distance from the mixing point. The observed SAXS signal decay (I(q) ∝ q^−2^) confirms that the measured droplets had radii larger than 300 nm.

#### Data fitting with Gaussian blobs

To estimate the mass distribution on the length scale of many Bik1 dimers, we fit the SAXS data using 4000 Gaussian blobs with a standard deviation σ of 40 Å. The blobs are arranged in a spherical volume with a diameter of 10’000 Å, which is large enough to cover the largest length scale that can be resolved by our SAXS measurements. The use of Gaussian blobs for our data fitting approach is convenient as the real-space and the reciprocal space representations are described by Gaussians. This approach provides a possible configuration that is orientationally averaged for the fit to the SAXS data, as many droplets contributed to a measurement. Therefore, the scattering curve for the fit is calculated as the average resulting from 300 random orientations of the momentum transfer vector. Least-square fits to the SAXS data were performed using the self-developed code written in the Julia language.^62^

The form factor of the sphere containing the Gaussian blobs must be suppressed, as it would disturb the fit at low *q*. This is achieved by using a Gaussian window placed at the center of mass of all blobs used to fit the measurement and decaying from the origin to the outward. The scattering amplitudes of all blobs in the volume are multiplied with this window function, which corresponds to a convolution with a Gaussian in reciprocal space. The window function is used with a standard deviation σw= 3000 Å, which sets the *q* resolution of the fit to 2π/σw ≈ 0.002 Å^−1^.

#### Data fitting with spherocylinders

Our SEC-SAXS results revealed an elongated, cylindrical conformation for the Bik1 dimer, which is considerably different than Gaussian blobs. They further suggested that the overall conformation of the protein is dominated by its two-stranded coiled coil with little, if any, contributions from its N- and C-terminal domains. Based on our computational analysis, this coiled coil can be divided into three segments that are connected by flexible linkers. To fit our SAXS data with a more realistic representation of the Bik1 dimer, we thus constructed three models with increasing complexity: model 1 consists of a single spherocylinder with a radius of 12 Å and a length of ≈ 300 Å; model 2 consists of two linked spherocylinders with radii of 12 Å and lengths of 155 Å and 135 Å, respectively; model 3 consists of three linked spherocylinders with radii of 12 Å and lengths of 155 Å, 76 Å, and 50 Å, respectively.

To fit the SAXS data using the three models, we placed 50’000 or 100’000 model copies according to the mass distribution obtained from the Gaussian-blob fit in a spherical volume with a diameter of 6000 Å. This chosen diameter restricts our analysis to *q* > 0.004 Å^−1^ or a maximum distance of about 1500 Å, which is much larger than the length of a fully elongated Bik1 dimer and is therefore acceptable. Due to the reduced volume, the window function is used with σw = 1500 Å and, therefore, the *q* resolution of the fit is reduced to 2π/σw ≈ 0.004 Å^−1^, which does not limit our analysis of the SAXS data. All least-square fits to the SAXS data were performed using Julia code^62^ for the calculation of the scattering signals resulting from spherocylinders as well as for the detection of spherocylinder clashes in the volume used to fit measurements.

For fits using model 1, spherocylinders with random orientations are placed close to the centers of the Gaussian blobs before translations and reorientations of them are carried out to improve the fit to the SAXS data. While translations and reorientations are chosen randomly, only those improving χ^2^ of the fit are accepted, while all others are rejected. Good fits were obtained for all three SAXS measurements. This result was achieved with two volume fractions, φ ≈ 0.05 and φ ≈ 0.1, for all three SAXS measurements. Model 2 approximates a Bik1 dimer with two spherocylinders with a flexible angle α between the cylinder axes. In the stretched-out configuration (α=180°), model 2 mimics model 1, while the two spherocylinder components lie parallel to each other at α=0° in the fully folded-back configuration. The result obtained with model 1 was used to start the data fitting using model 2. The longer of the two segments of model 2 was superimposed on one end of each copy of model 1, and the second segment was added with a random orientation in a manner to avoid clashes with neighboring model copies. The second segment was placed by ‘rolling’ its spherical cap on one of the first segment until the desired orientation was reached. The fit to the SAXS data was performed with random translations of model 2 segment pairs and random reorientations of the first or second segment of the pair, always maintaining the link of the two segments and avoiding clashes with neighboring model copies. Analogous to the transition from model 1 to model 2, the data fitting with model 3 was started using the result obtained with model 2. Good fits to the data were obtained, and the orientation of the three model segments was close to random with a slight tendency towards a fully stretched-out configuration for both angles α and *β* between model segments (for angle distributions, see **Figure S9D-F**). For the minimization of χ^2^, the same procedure using reorientations and translations is used for all three models. The fits to the data obtained with models 1, 2, and 3 were of comparable high quality.

The local nematic order parameter, S, was determined from the model 1 fits obtained with 50’000 or 100’000 model-1 copies. The model-1 copies in cubic boxes with side length 400 Å were considered to calculate the local order parameter tensor. The local order parameter is obtained as *S* = 3/2\**e*, where *e* is the largest positive eigenvalue of the order parameter tensor (**Fig. S10**). With boxes containing 30 or more model-1 copies, the order parameter is below 0.4, which we interpret as an orientationally unordered configuration. With 50’000 copies, the order parameter can be higher than 0.4 when *N*<30 and the spherocylinders happen to be quite well aligned.

## Supporting information

Bik1_movies

## Data availability

Custom R script code and MS files are available in the PRIDE archive, a public data repository for MS proteomics data (https://www.ebi.ac.uk/pride/), with the following IDs: PXD050928, PXD050929, PXD050930, PXD050931. The SEC-SAXS datasets can be found in the SASDB archive, Small Angle Scattering Biological Data Bank (https://www.sasbdb.org/) with the following accession codes: SASDUT6, SASDUU6, SASDUV6. The SAXS datasets collected for the phase-separated droplets of Bik1 are deposited in the Zenodo (https://zenodo.org/) repository, with the following DOI: 10.5281/zenodo.10944717.

## Acknowledgments

We thank S. Meier for the purification of mNG-Bik1 FL and the staff of the DLS beamline B21 of the Diamond Light Source for the provision of and excellent assistance with SAXS beamtime. We are grateful to A. Leitner for his insights into cross-linking strategies, for the access to the data analysis platform, and the use of chromatographic devices. We acknowledge financial support from the Swiss National Science Foundation Sinergia grant CRSII5_189940 (to E.R.D. and M.O.S.). M.P.C. has received funding from the European Union’s Horizon 2020 research and innovation program under the Marie Skłodowska-Curie grant agreement No 884104 (PSI-FELLOW-III-3i). This work benefited from access to Beamline B21 at Diamond Light Source and has been supported by iNEXT-Discovery, project number 871037, funded by the Horizon 2020 program of the European Commission.

## Author contributions

M.P.C. and M.O.S. designed the project and wrote the first draft of the manuscript. M.P.C. and T.G. purified proteins and performed functional studies. M.P.C. and F.U. designed, performed, and analyzed crosslinking experiments. K.R. performed and analyzed confocal microscopy experiments. M.P.C. and I.V. prepared and analyzed SEC-SAXS measurements. M.P.C., T.G., C.P., A.M., and U.G. designed and performed SAXS measurements of phase-separated Bik1 samples, and U.G. analyzed and modeled the obtained results. M.O.S. supervised the entire study. All authors contributed to the reviewing and scientific interpretation of the manuscript.

