## Supplemental for "Phase separation of a microtubule plus-end tracking protein into a fluid fractal network"

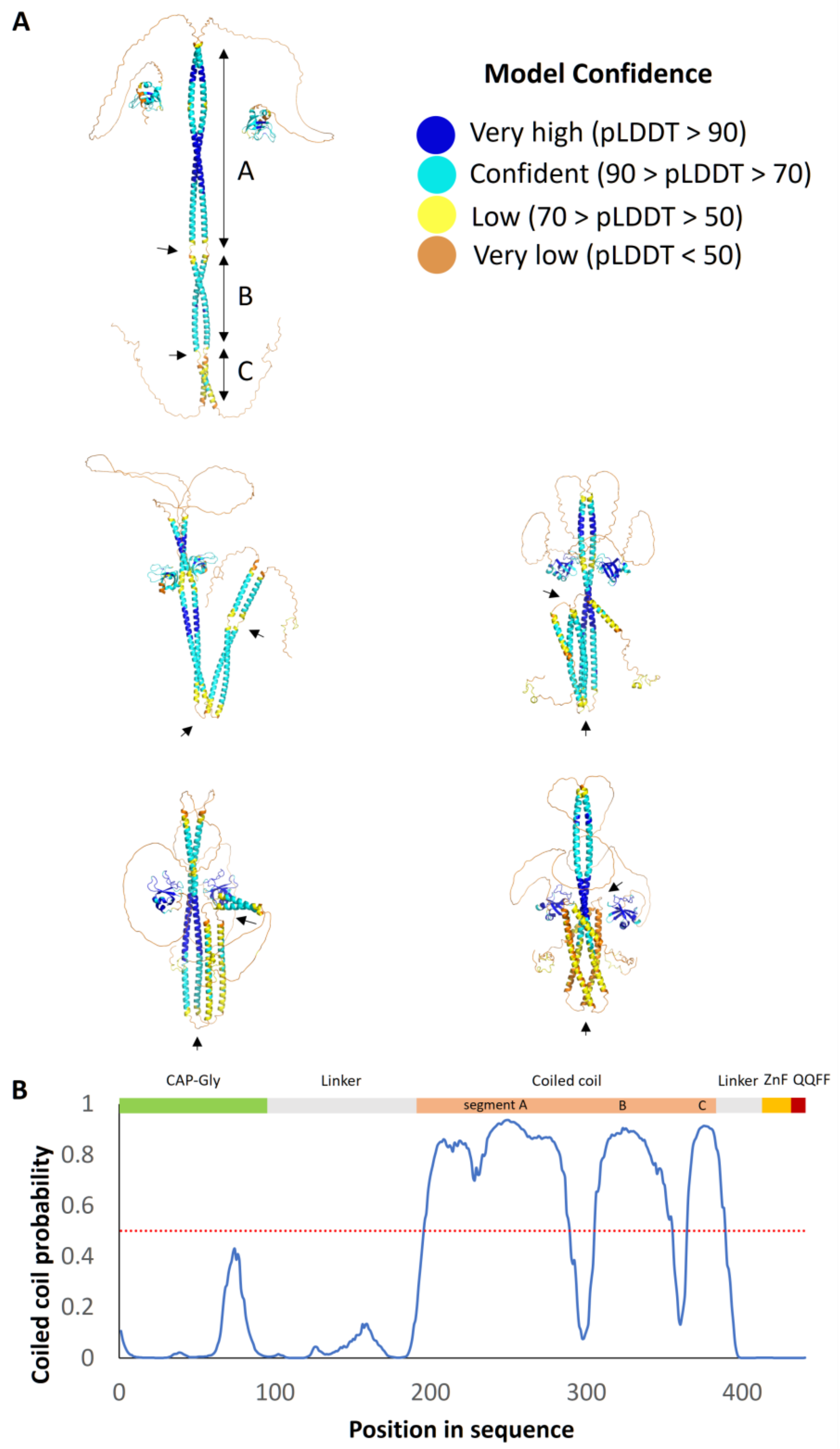


**Figure S1. Computational predictions of the Bik1 structure.** A) Different possible structures of the Bik1 dimer predicted by AlphaFold.[1] Structures are color-coded according to the pLDDT score: regions with a high pLDDT score (>90) are expected to be modeled with high accuracy, while regions with a low pLDDT score (<70) were modeled with low confidence. Black single arrows indicate predicted discontinuities in the coiled-coil domain; double black arrows indicate the three coiled-coil segments A, B, and C. B) Sequence-based coiled coil prediction of Bik1 by DeepCoil.[2] The blue line represents the probability of coiled-coil formation per residue; the red dashed line indicates the 50% probability threshold. ZnF, zinc finger domain; EEY/F-like motif Gln-Gln-Phe-Phe.

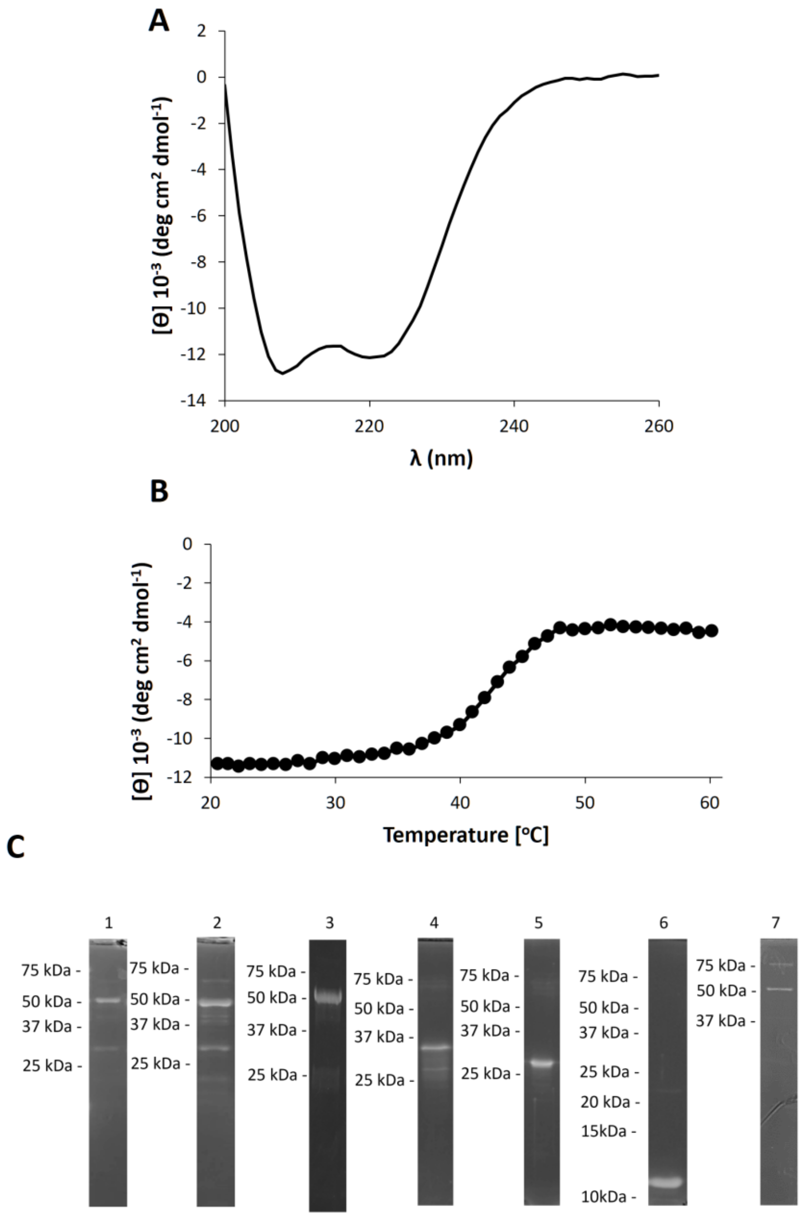


**Figure S2. Quality assessment of Bik1 samples.** Spectrum (A) and thermal unfolding profile (B) recorded by CD at 222 nm for Bik1 FL in high-salt buffer conditions (20 mM Tris-HCl, pH 7.4, supplemented with 500 mM NaCl and 10% glycerol). C) Coomassie Blue-stained SDS-PAGE analysis of Bik1 protein variants used in this study: 1, Bik1 FL; 2, Bik1 FL cleaved; 3, Bik1 ΔQQFF; 4, Bik1 ΔCG; 5, Bik1 CC; 6, Bik1 CG; 7, Bik1 Δtail. Bik1 FL cleaved stands for the full-length protein that was incubated with PreScission Protease to cleave the N-terminal His-tag; the His-tag was removed by passing the cleaved sample through a Histrap column. All other samples contained an N-terminal His-tag.

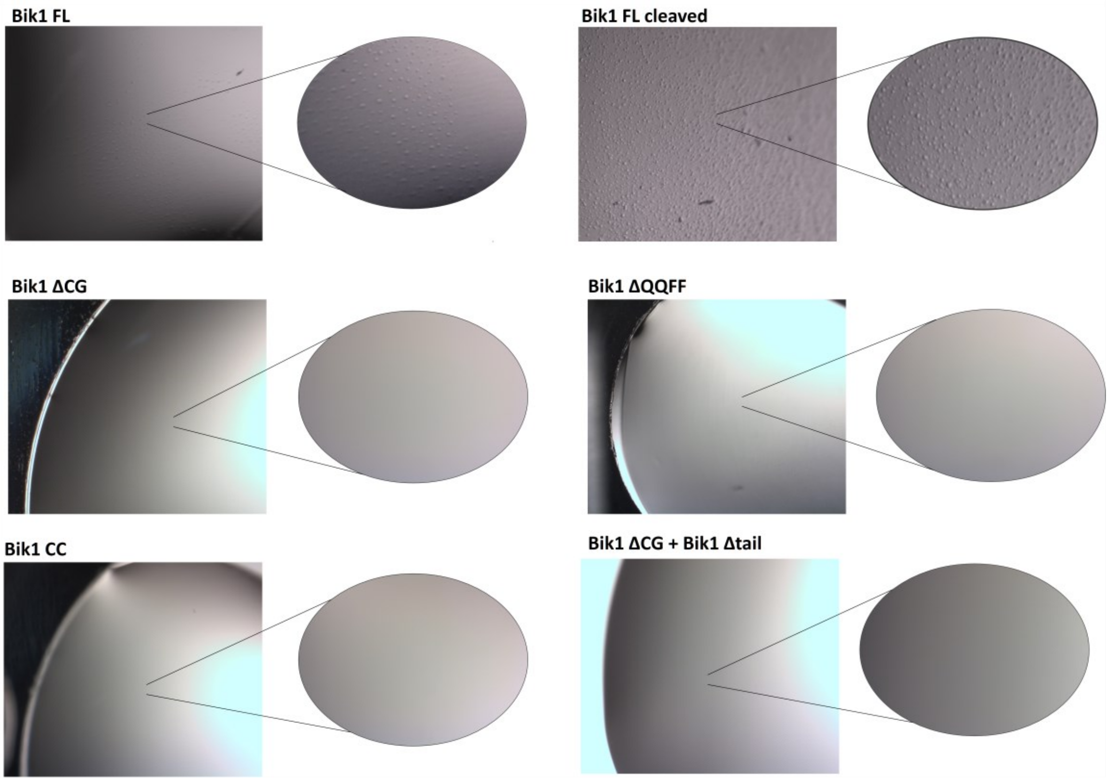


**Figure S3. Phase separation properties of N- and/or C-terminal Bik1 truncation mutants.** Light microscopy images of Bik1 FL and its truncated variants in low salt buffer conditions (10 mM Tris-HCl, pH 7.4, supplemented with 250 mM NaCl, 0.5 mM DTT, and 5% glycerol). Bik1 FL cleaved stands for the full-length protein that was incubated with PreScission Protease to cleave the N-terminal His-tag; the His-tag was removed by passing the cleaved sample through a Histrap column. Despite high protein concentrations (50-100 µM), Bik1 variants containing truncations in N- and/or C-terminal regions were not observed to undergo phase separation under the conditions applied.

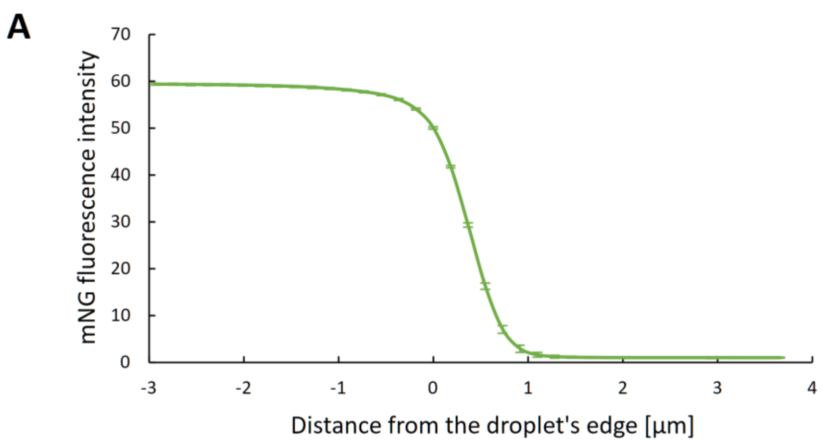


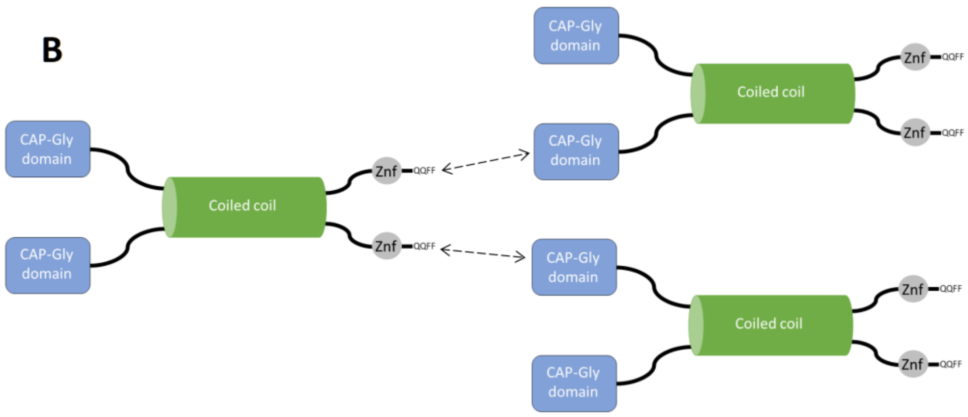


**Figure S4. Analysis of phase separated mNG-Bik1 FL droplets and Bik1 intermolecular interaction scheme.** A) The average intensity of mNG throughout mNG-Bik1 FL droplets (**Figure 1E**). B) Possible intermolecular interactions between Bik1 dimers mediated by the protein's N-terminal CAP-Gly and C-terminal tail domain. Unstructured regions of the protein are shown as black solid lines. ZnF, zinc finger domain.

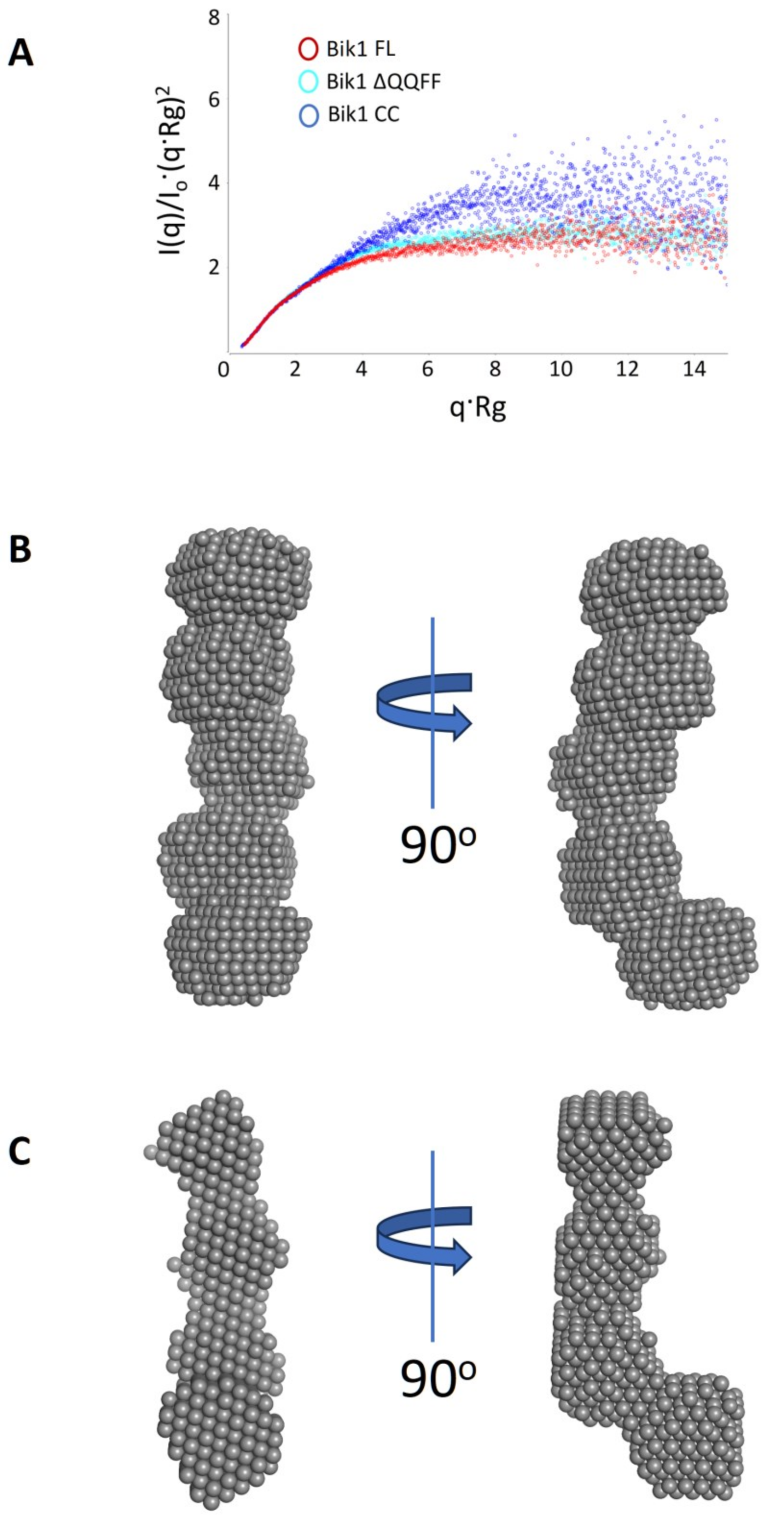


**Figure S5. SEC-SAXS analysis of Bik1 variants.** A) Dimensionless Kratky plots obtained for Bik1 FL, Bik1 ΔQQFF, and Bik1 CC. B and C) DAMMIF generated molecular envelopes (beads model) of Bik1 ΔQQFF (B) and Bik1 CC (C) calculated by averaging 19 models (out of a total of 20) resulting in NSD values of 0.71 ± 0.05 and 0.61 ± 0.04, respectively. The resolution of ensembles is 65 ± 5 Å for Bik1 ΔQQFF and 43 ± 3 Å for Bik1 CC.

**Table S1. $R_g$ and $D_{max}$ values calculated from SEC-SAXS data.**

| Bik1 variant | Protein concentration [mg/ml][1] | $R_g$ [Å] | $D_{max}$ [Å] |
|---|---|---|---|
| Bik1 FL | 8 | 98 | 320 |
| Bik1 FL | 6 | 91 | 280 |
| Bik1 FL | 4 | 90 | 280 |
| Bik1 ΔQQFF | 8 | 97 | 320 |
| Bik1 ΔQQFF | 6 | 94 | 294 |
| Bik1 ΔQQFF | 4 | 94 | 300 |
| Bik1 CC | 8 | 83 | 298 |
| Bik1 CC | 6 | 80 | 270 |
| Bik1 CC | 4 | 79 | 250 |

[1]Protein concentrations were measured before the protein was injected into the SEC column.

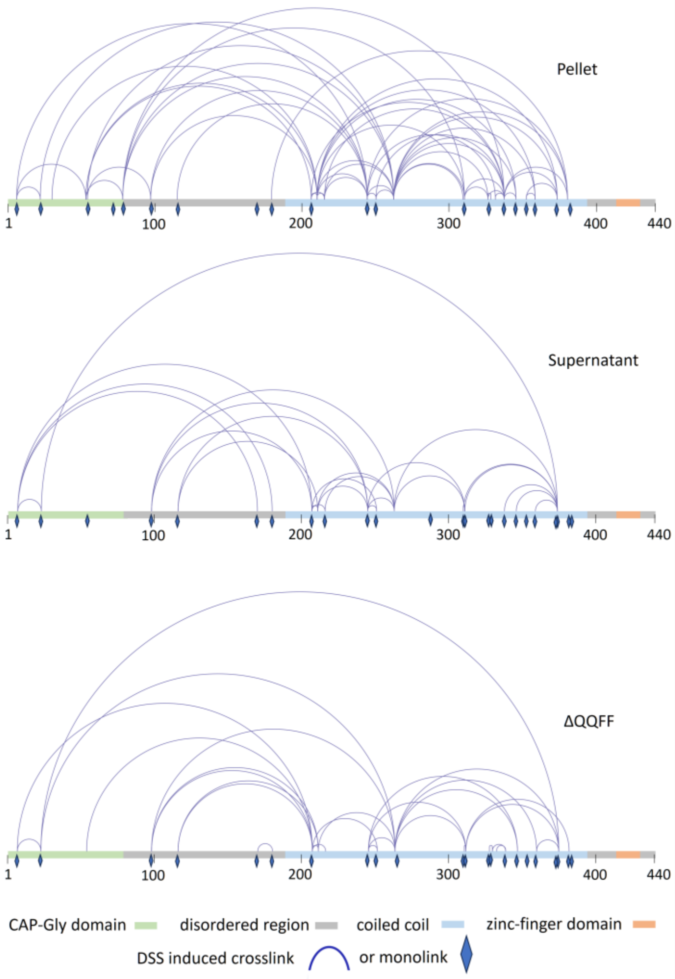


**Figure S6. XL-MS experiments.** Crosslinks and monolinks identified using the DSS crosslinking reagent in the pellet and supernatant of Bik1 FL and Bik1 ΔQQFF. Crosslinks linking different regions of the proteins are depicted as blue lines; monolinks are shown as blue rhombi. Protein domains are color-coded as follows: CAP-Gly domain, light green; disordered region, gray; coiled coil, light blue; zinc-finger domain, beige.

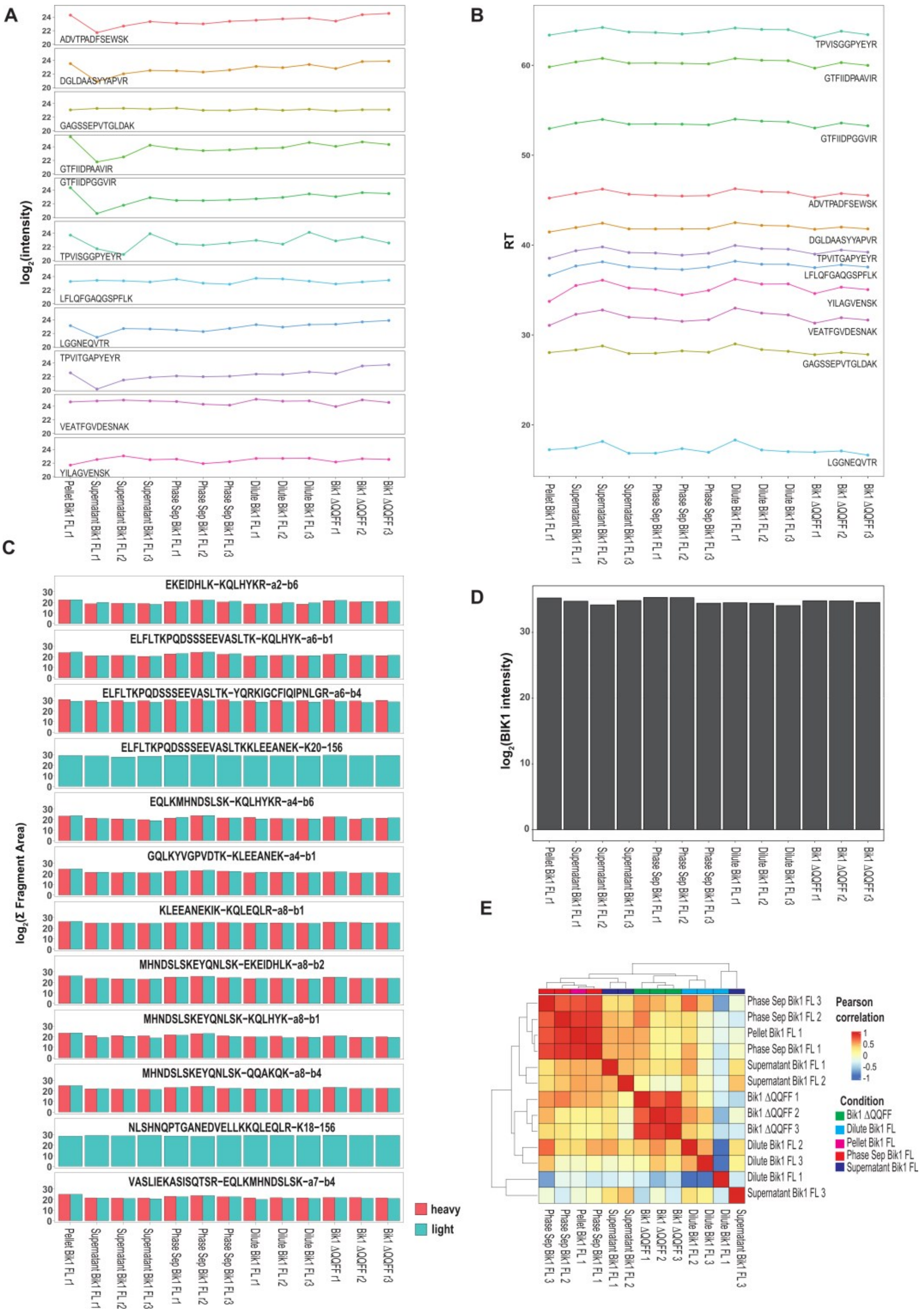


**Figure S7. Quality control for targeted XL-MS analysis.** A) Quality control to probe the sensitivity of external reference peptides (iRT) during the targeted analysis acquisition. The peptide abundance (log2 of the sum of the top five transitions) is provided for all acquired runs. B) Quality control to probe the retention time of external reference peptides (iRT) during the targeted analysis acquisition. The retention time is provided for all acquired runs. C) Intensity abundance (expressed as log2) for crosslinked peptides monitored across all acquired runs. Cyan and red bars indicate the abundance of heavy and light crosslinked peptides, respectively. D) Intensity abundance of Bik1 FL (expressed as log2) for all acquired runs. The reported intensity is derived from the average intensity of the two Bik1 peptides MVLEEVQPTFDR and HSGNQQSMDQEASD-HHQQQEFGYDNR. Bik1 abundance serves as a normalization factor for crosslink intensity to control protein recovery levels. E) Unsupervised hierarchical cluster based on Pearson correlation for crosslinked peptides identified.

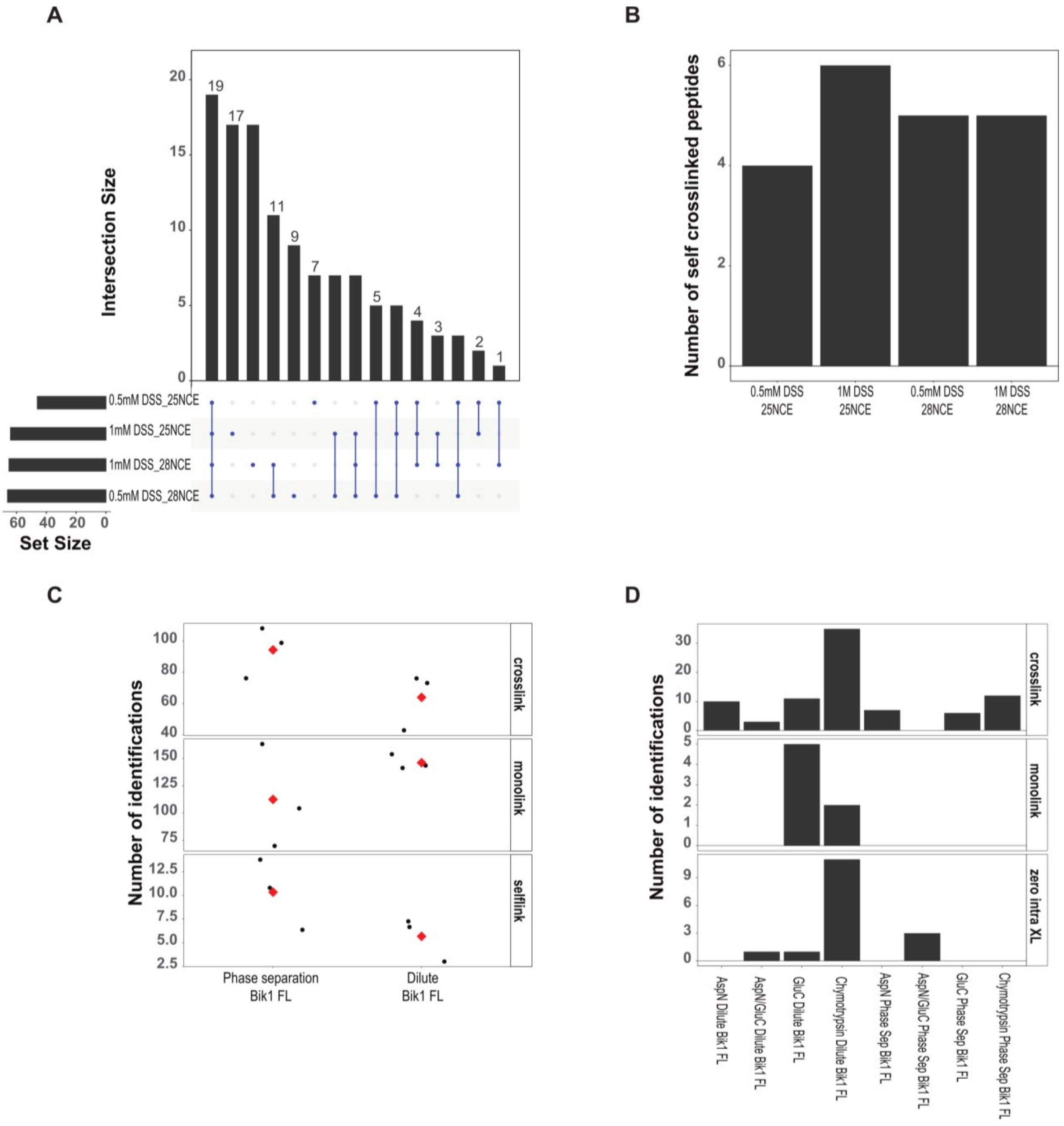


**Figure S8. Analysis of different crosslinked Bik1 FL samples.** A) Upset plot of identified crosslinked peptides using different concentrations of DSS and distinct acquisition method based on varying collision energy. B) Identification of self-linked peptides at different experimental conditions (different concentrations of DSS reagent and collision energy) C) Quantification of crosslinked peptides (intra-crosslinks, monolinks, and self-link) using DSS crosslinking reaction in both phase-separated and dilute conditions. Each data point represents technical replicates; red diamonds indicate the average value. D) Quantification of crosslinked peptides (intra-crosslinks, monolinks, and zero link crosslink) using PDH crosslinking reaction and different proteolysis mix (AspN; AspN/GluC; GluC; chymotrypsin) in phase separated and dilute Bik1 FL samples.

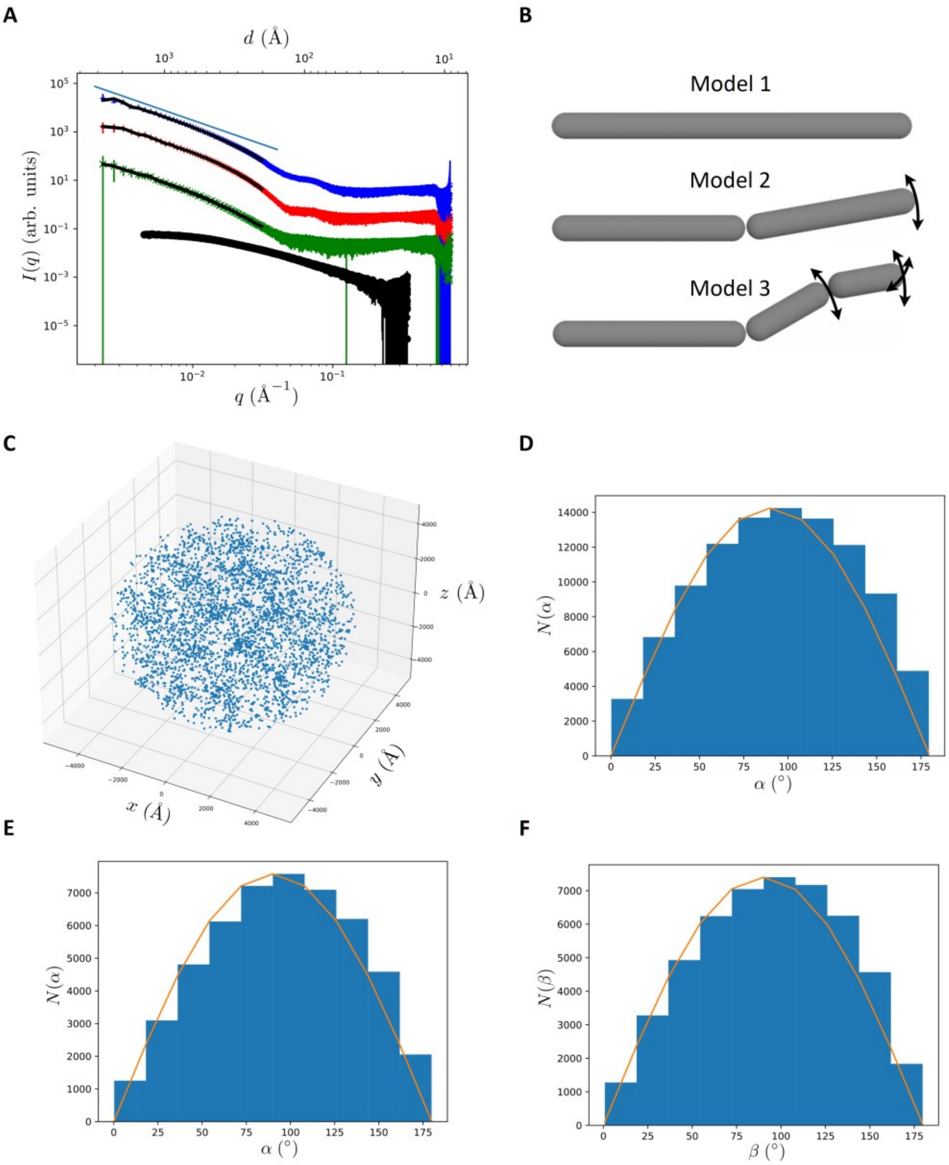


**Figure S9. SAXS analysis of phase separated Bik1.** A) SAXS data of phase-separated Bik1 FL measured in triplicates (blue, red, and green curves) and fits (black curves) obtained with 4000 Gaussian blobs in the *q* range (0.002 Å$^{-1}$ - 0.03 Å$^{-1}$) corresponding to distances larger than a fully extended Bik1 dimer. The SAXS data have been shifted with respect to each other in the direction of the Y-axis by one decade for clarity. The form factor obtained from our SEC-SAXS measurements of Bik1 FL (black curve) is shown for comparison. The straight blue line above the SAXS data represents $I(q) \sim q^{-2}$ behavior expected for a fractal network with fractal dimension equal to 2. B) Schematic representations of Bik1 models 1,2, 3 that were used for data fitting. C) Heterogeneous configuration of Gaussian blob positions in the spherical volume with a diameter of 1 µm resulting from a fit shown in panel A. Distribution of α angle in the model 2 (D) and model 3 (E). F) Distribution of β angle in model 3.

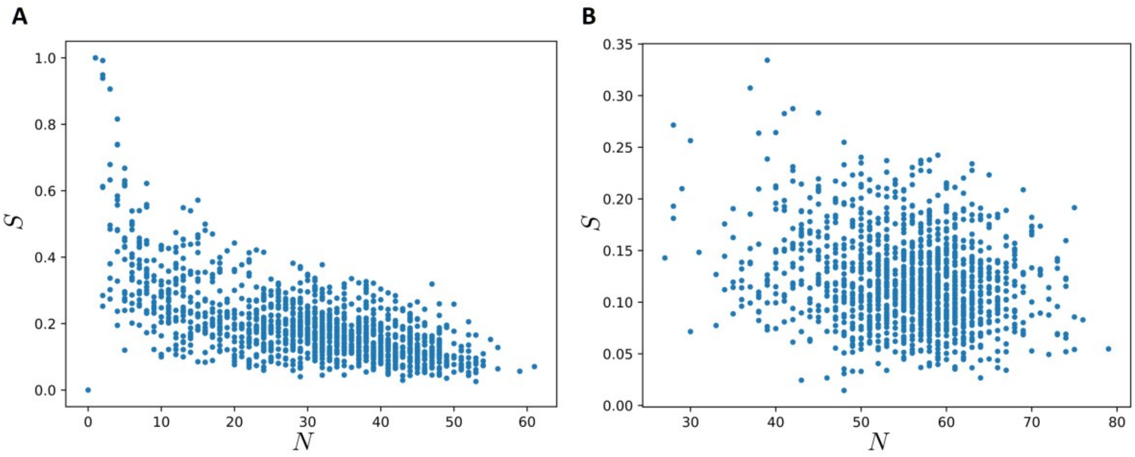


**Figure S10. Local nematic order parameter S versus number of model 1 copies, N.** The order parameter is determined locally in cubic boxes with side length 400 Å with respect to the average director determined as the average orientation of the model 1 copies in the box. (A) 50’000 model 1 copies and (B) 100’000 model 1 copies used in the fit to the SAXS data.
